# Tree crowns as meeting points of diversity generating mechanisms – a test with epiphytic lichens in a temperate forest

**DOI:** 10.1101/2020.01.03.894303

**Authors:** R. Patzak, R. Richter, R.A. Engelmann, C. Wirth

## Abstract

Forest canopies are hotspots of biodiversity even in temperate forests but which and how many ecological mechanisms contribute to the high diversity remains elusive. This biodiversity is not distributed evenly throughout the complex fractal structures formed by individual tree crowns. They are non-stationary, constantly expose new surface habitat via growth, and create contrasting abiotic conditions. These features give rise to a range of vertical gradients in habitat optimality, heterogeneity, available surface area and time for succession - all known to be mechanisms shaping diversity patterns. Using a canopy crane facility and epiphytic lichens on Fraxinus excelsior and Quercus robur as model system, we aim to assess the relative importance as well as the interplay of these mechanisms in shaping biodiversity patterns within tree canopies by detecting their distinct mechanistic fingerprints. Lichen species richness exhibited a hump-shaped vertical pattern, skewed towards the top of the crown. This pattern was observable at both the level of individual plots and that of aggregate height layers and it was correlated with lichen cover. Also, a vertical gradient in species composition was found and could be related to species traits known to reflect successional niches such as dispersal mode and growth form. Habitat heterogeneity and available surface area have been found to have little effect on vertical lichen diversity patterns. We conclude that the vertical lichen diversity patterns in the tree crown are mainly shaped by the successional accumulation of species along a branch age gradient and a pronounced vertical gradient in environmental optimality from harshly exposed young branches at the top crown over suitable habitats with a balance in light and humidity towards the limiting light conditions in the dim understory. At the level of the whole canopy, successional and environmental niche dynamics jointly operate to generate lichen diversity.

## 1 Introduction

Forest canopies are known as hotspots of biological diversity (Ozanne et al. 2003, Nakamura et al. 2017). Approximately 28.600 vascular plant species inhabit tree crowns as epiphytes, which comprises about 10 % of the total vascular flora (Nadkarni 1994). Extrapolations based on host specificity of arthropods with respect to tree species, resulted in a total number of 6.1 million terrestrial arthropod species harbored in the tropics (Hamilton et al. 2010). This high diversity of arboreal organisms has been attributed to a number of unique features of tree crowns including the richness of substrates (Barkman 1958, Fritz 2009) and the availability of light energy in comparison with the forest floor. Furthermore, pronounced environmental gradients spanning from dark moist habitats in the lower to sunlit dry habitats in the upper canopy layer (Parker and Brown 2000) create a wealth of niches for canopy organisms. Next to a vertical niche gradient, the complex architecture of tree crowns also creates horizontal variation in environmental conditions (McCune et al. 2000). The extent of this variation is expected to change systematically with height (Parker and Brown 2000). In addition, and thus far not considered in the literature, the availability of branch surface area increases with height as the branching process divides a given wood volume into successively thinner branches with a higher surface to volume ratio. Habitat area for bark-dwelling organisms thus increases with height. According to positive species-area relationships (Connor and McCoy 2000) this alone should allow for higher diversity in the top layer of tree canopies. Using a crane facility, we developed a novel sampling design for canopy research with the goal to quantify the relative importance of different diversity generating mechanisms of epiphytic diversity in tree crowns using lichens as model organisms.

Epiphytic lichens represent a ubiquitous component of canopy communities. Even in temperate forests the species richness of lichens may exceed that of forest trees by at least an order of magnitude (Ellis 2012). Here we employ arboreal lichen species as model organisms to explore the relative importance of mechanisms shaping patterns of biodiversity in tree crowns of two temperate tree species (*Fraxinus excelsior* L. and *Quercus robur* L.). We sampled lichens species richness (α diversity) and abundance in subplots positioned in five canopy layers and at the trunk. The number of plots in these layers was proportional to the available bark surface area. The total number of species per layer is referred to as γ diversity, their turnover between plots within a layer as β diversity.

For autotrophic organisms like lichens, radiation is inarguably a key resource and influences growth rates (Hilmo 2002). During leaf-on, the amount of radiation transmitted into the canopy decreases quickly from a shallow zone of bright light conditions at the top of the canopy to darker regimes in deeper canopy layers. The bottommost zone around the trunk receives only about 5 % of the radiation above the canopy in temperate deciduous forests (Parker 1997). However, in poikilohydric lichens, photosynthesis may also be quickly limited by water availability (Sillett and Antoine 2004). Although lichens are able to endure severe periods of drought, in such periods they are not metabolically active (Palmqvist 2000, Kranner et al. 2014). With increasing height in the canopy, lichens are exposed to drying winds and high temperatures, while lower canopy layers provide more sheltered microhabitats. Hence, optimal conditions for lichen productivity and survival might be found in intermediate crown layers where the joint availability of light and moisture is highest (Figure). In this region, a higher productivity may sustain higher population densities. This may allow even rarer species to establish and persist (Wright 1983) which in turn should lead to a higher lichen diversity in this zone, both across the entire crown layer (γ diversity) and locally at the level of subplots (α diversity).

Orthogonal to vertical environmental gradients there is also horizontal variation in environmental conditions. Whereas the topmost layers are fully illuminated and well-coupled with the atmosphere and are thus uniformly bright, dry and exposed to wind (Unterseher and Tal 2006), the bottom layers tend to be uniformly dark and moist. However, in the transition zone the complex architecture of tree crowns with their clustered branches and foliage creates a patchy mosaic of microsites varying in microclimate (light, rain interception, wind exposure) and structure (branch sizes and inclinations, bark roughness) (McCune et al. 2000, Parker and Brown 2000). The resulting horizontal variation in substrate and microclimate in mid-canopy layers may promote the co-occurrence of specialists each with preferences for particular habitat types (Connor and McCoy 2000). As a result, the γ diversity of mid-canopy layers is expected to be high as a consequence of high species turnover between contrasting microsites (β diversity), but not because of peaking α diversity.

Another potentially important factor impacting composition and diversity of epiphytic communities are successional changes brought forth by the emergence of new habitat surface as trees grow in height and produce new branches. Colonization time on young branches in the top canopy is shorter than on old branches further down (Ellis 2012). Furthermore, during growth, lichen themselves change their microenvironment by altering branch surface structure, facilitating further establishment and increasing moisture interception (Pypker et al. 2006) or by producing allelopathic compounds as competitive means (Lawrey 1986). Thus, autogenic successional drivers may relate to branch age (Rogers 1988, Ellis and Coppins 2006, Johansson et al. 2007), although the continuous growth of the host tree coincides with drastic changes in microenvironment making allogenic succession more prevalent (Stone 1989). This leads to successional sequences of lichen community composition (Degelius 1964, 1978, Rogers 1988, Hilmo 1994, Wirth et al. 1999). Young top-layer branches may host lichen communities consisting of a limited set of fast colonizing early-successional species (Rogers 1990), while lichen communities on old branches, representing late stages of succession with high cover, may have lost species due to the exclusion by more competitive lichens and/or bryophytes (Armstrong and Welch 2007, Fritz 2009). As a consequence, lichen diversity is expected to increase from the top-layer downwards to regions where species of different successional stages co-occur before it decreases again (Degelius 1964, Hilmo 1994), thus creating a hump-shaped pattern, referred to as mid-succession peak (Johansson et al. 2007).

According to the species-area relationship (SAR), a landmark theory in ecology tested for many taxa, habitats and scales (Connor and McCoy 2000), vertical gradients in total available branch surface area may strongly affect vertical patterns of γ diversity. Although wood volume tapers with height to some extent, the power-law increase in surface-to-volume ratio in progressively thinner branches leads to a sharp increase of bark surface towards the top of the canopy. Thus, the major part of surface area available for epiphytes can be found in the upper canopy. According to SAR, this increase in available area towards the treetop should be paralleled by an increase in species richness. While an increase of lichen biomass with height has been reported (Hale 1952, Ellyson and Sillett 2003, Boch et al. 2013), we are not aware of any study considering SAR as potential driver of vertical patterns in lichen diversity in canopies let alone its component processes ‘area per se’, ‘habitat heterogeneity’ and the ‘passive sampling process’ (Connor & McCoy, 2000). With the SAR mechanism, γ diversity is expected to increase monotonously with height.

This study aims to disentangle the effects of mechanisms generating patterns of lichen diversity in tree canopies. With the exception of the SAR mechanism, all other mechanisms introduced above are hypothesized to produce a mid-canopy peak of γ diversity at the level of canopy layers (Figure) and are thus indistinguishable without additional information. However, each of the four mechanisms is assumed to produce a characteristic fingerprint with respect to patterns of diversity components such as α and β diversity, gradients of trait expression and relationships to underlying patterns of niche predictors (see Table 1 in the method section for a summary of our predictions). It is important to note that the four mechanisms are not mutually exclusive. Here, we present a sampling design, theoretical framework and analysis scheme that allows us to holistically assess the relative contributions of the four mechanisms generating patterns of lichen diversity on two tree species of a Central European floodplain forest. To this end, we employ a variety of tools such as linear models, variance partitioning, null model comparisons and structural equation models, where processes and their underlying mechanistic hypotheses are represented by specific pathways.

**Table 1:**
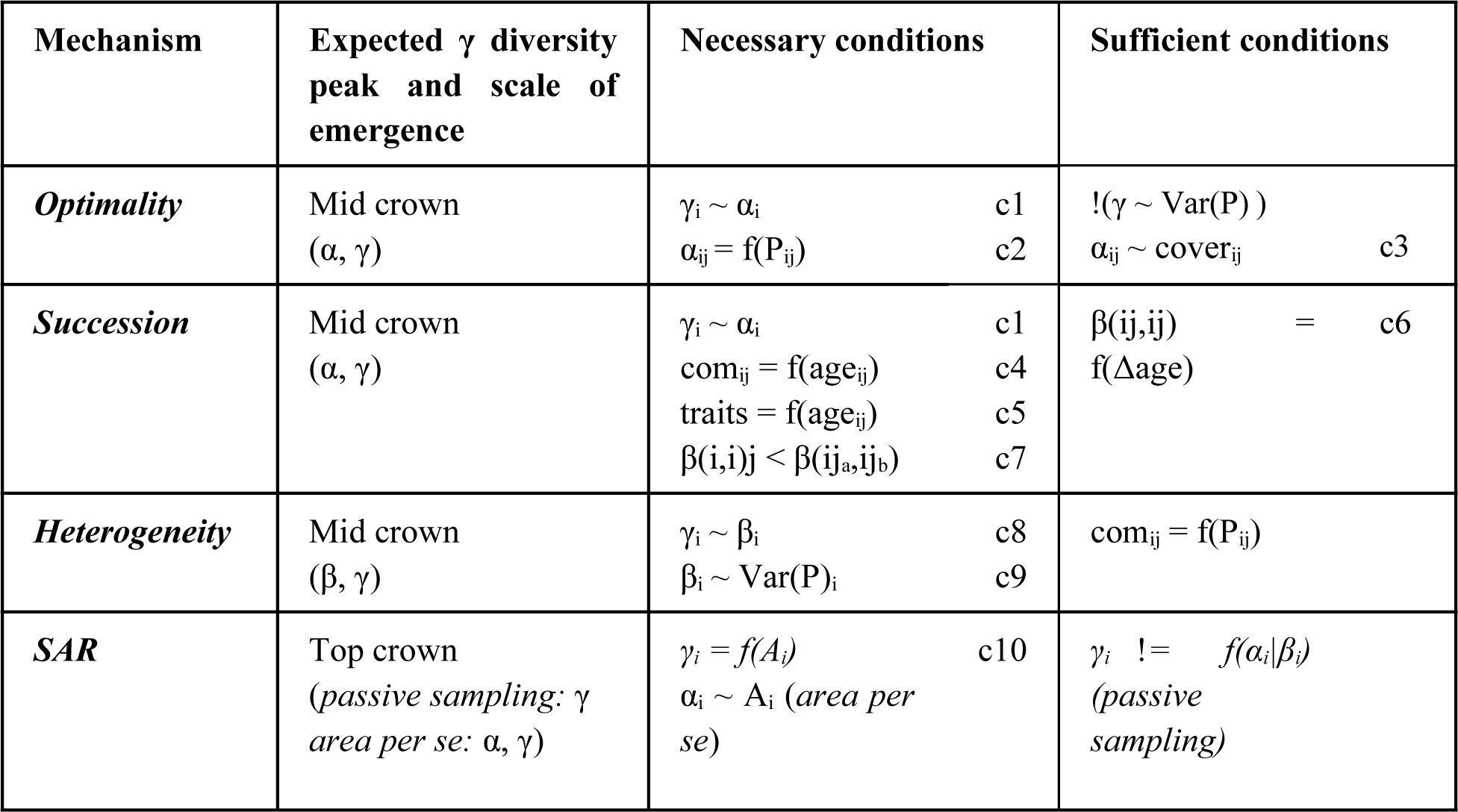
Evidence matrix: Expected observations according to each hypothesized mechanism. The first column details were in the crown a peak in diversity is to be expected and which measure or scale of biodiversity will be affected. Here, ‘mid crown’ does not necessarily refer to the mathematical midpoint of the crown but rather may be slanted to either top or bottom of the canopy. The second and third column feature conditions that have to be observed necessarily or are indicative of a mechanism, respectively. Symbols and abbreviations used are: α, β, γ = α, β or γ diversity; com = community composition; A = branch surface area; P = habitat predictors (e.g. branch inclination, LAI); age = branch age, ‘traits’ is referring to a successional species trait gradient; i = layer marker, j = plot marker (j_a_ and j_b_ mark different layers). The numbers following the conditions in the third and fourth column are for reference in *Figure 2*.

We expect to find evidence that the several mechanisms operate in *parallel* and mutually reinforce each other to produce a distinct mid-canopy peak of γ diversity at the level of canopy layers. We further expect that this diversity peak is displaced upwards as a consequence of the SAR ‘passive sampling’ mechanism operating on the increasing availability of substrate area with height. We hope to contribute to an understanding of tree canopies as meetings points of diversity shaping mechanisms that could explain why tree canopies are often found to be hotspots of biodiversity.

## 2 Methods

### 2.1 Study site and tree selection

The study was conducted at the Leipzig Canopy Crane facility (LCC). The crane site is located within the nature reserve “Burgaue” in the northwestern part of a floodplain forest that traverses the city of Leipzig in Saxony, Germany (51°20’16” N, 12°22’26” E). Leipzig lies in the transient area of maritime to continental climate with mean annual rainfall of 556.9 mm and a mean temperature of 9.8 °C in average (Kaiser 2014). The crane is a Liebherr 71 EC tower crane with a height of 40 m and horizontal reach of 45 m, which is mounted on a 120 m railroad track. Thus, it provides easy access to the forest canopy within an area of 1.65 ha. At the time of sampling this area comprised ∼800 trees with diameter at breast height (DBH) > 5 cm belonging to 17 species (10/ha), with *Acer pseudoplatanus* L.*, Fraxinus excelsior* L. and *Tilia cordata* Mill. being the most dominant tree species*. Fraxinus excelsior* and a few exceptionally large individuals of *Quercus robur* L. represent the individuals with the highest basal area. Of these species, five large individuals each were chosen as sampling targets.

### 2.2 Sampling design

To enable us to account for area-specific effects on lichen diversity patterns, each tree was subdivided into five equally spaced crown layers and a sixth trunk layer. The crown base was defined as the point of separation from the main trunk into erect branches having contact to the upper crown region. Lacking such contact, a branch was assigned to the trunk layer. Within each layer, sampling plots were randomly placed with a minimum distance of 1 m to each other. Their number was proportional to the share of total bark surface of the respective specific layer, whereby the minimum number of plots in the lowermost crown layer was set to five. The available surface area within each layer was approximated by means of a Dirichlet regression model (Maier 2014), trained on a dataset containing biometric data collected via *random branch sampling* (Gaffrey and Saborowski 1999) originating from 385 single broadleaf trees (Riedel and Kändler 2017). Different models were used for modelling surface area proportions for each phorophyte species (S1). The *F. excelsior* model was using diameter at breast height (DBH) as predictor for a second-order polynomial model, the *Q. robur* model used a logarithmic model with DBH as predictor and the ratio crown base height / total tree height as additional predictor. This resulted in a total number of 55 - 99 plots per tree. Each sampling plot consisted of two sampling areas with dimensions of 1.5 x 33.3 cm (100 cm^2^ in total) on the upward and downward facing side of a branch. The dimensions have been chosen to accommodate vastly differing branch diameters (range: >1 – 87 cm) without changing the shape of the sampling area. On the trunk and other vertical branches, the sampling areas were placed on opposite cardinal directions alternating between north/south and east/west. Sampling was carried out from October 2016 to July 2017. For each lichen species, cover was estimated from occurrences using a 5 mm grid. Species were identified in the field if possible. Otherwise, voucher specimens were taken for later identification. Identification and naming convention followed Wirth, Hauck, & Schultz (2013). Some individuals of *Physcia adscendens* and *P. tenella* could not be determined at the species level due to insufficiently developed soralia and were thus grouped under *Physcia sp*. Note: this species aggregate does not include other species of the same genus, like *P. stellaris*. In two cases identification completely failed. These species were noted by their description (for example “sterile grey crustose lichen”). As additional parameters for each sampled branch the height above ground, diameter, inclination and distance and azimuth to the trunk was recorded using measuring tapes or calipers, an inclinometer, a laser rangefinder VERTEX VL 5 (Haglöf, Järfälla, Sweden) and a compass respectively. Branch age was modelled from branch diameter and species using a data set of branch cross sections collected off fallen trees in the area (S2). Bark roughness was estimated categorically according to an ordinal scale ranging from 1 (smooth) to 6 (rough). Bark lesions and deadwood were noted. Leaf area index (LAI) was measured at the plot level in August and September 2018 using a Plant Canopy Analyzer LAI-2200C (LI-COR, Lincoln, NE, U.S.A.).

### 2.3 Tracing the mechanistic pathways

The following hypothesized mechanisms are tested for their influence on driving epiphytic lichen diversity in the tree crown: (i) *Habitat optimality*, (ii) *Succession*, (iii) *Habitat Heterogeneity* and (iv) *Species Area Relations (SAR).* We propose that each of the four mechanisms produces a characteristic fingerprint with respect to the height gradients and correlations patterns of its diversity components (α, β, γ), gradients of trait expression and relationships to underlying patterns of niche predictors, which are summarized in Table 1. These mechanisms are not mutually exclusive, but presumed to interfere and interact with each other.

For the *Optimality mechanism* to operate, a mid-canopy γ diversity peak has to occur which is driven by a peaking α diversity, i.e. γ and mean α diversity are correlated at the canopy-layer level. It is further presumed that α diversity is related to one or several relevant environmental predictors. The optimal conditions should also be reflected by a higher cover of lichens. Under the *Optimality mechanism* one would not expect a positive relationship between the within-layer variability of environmental variables and γ diversity and no systematic species turnover along the vertical canopy gradient.

The *Succession mechanism* is expected to produce a mid-canopy γ diversity peak driven by a peaking α diversity as above. It is an additional necessary condition that that there is a systematic species turnover along the gradient of branch age, as a proxy of successional time. There also needs to be systematic species turnover along the vertical canopy gradient, as mean successional time decreases with height, making plots within a layer more similar to each other than plots in other layers. Consistent with the *Succession mechanism* is a positive correlation of within-layer β diversity and the variation in branch age.

The *Habitat Heterogeneity mechanism* is expected to produce a mid-canopy γ diversity peak that is driven by a peaking β diversity, but not α diversity as in the first two mechanisms mentioned above. As α and β diversity are often correlated (Chase et al. 2011), this implies that observed dissimilarities between layers have to exceed the level of dissimilarity expected by a baseline α-β correlation. It is a necessary condition that β diversity is positively correlated with variation in habitat variables (inclination and bark roughness, LAI) pointing to niche diversity within layers. For the *SAR mechanism* to operate according to the *passive sampling* hypothesis *sensu* Connor & McCoy (1979), where a higher surface area acts as larger sample for colonists, γ diversity should correlate with available substrate area per layer and thus should peak in the top canopy layer. γ diversity can be independent of either α or β diversity. If available area is a strong determinant for species diversity, its peak may be skewed towards the top of the crown. In contrast, the *Optimality*, *Succession* and *Heterogeneity mechanisms* all may predict a richness peak towards the mid crown, as this region may offer a more balanced mix of sheltered, yet light habitats (McCune et al. 2000, Parker and Brown 2000), a transition zone from early to late successional species (Rogers 1988, Hilmo 1994) and environmentally and architecturally more diverse habitats respectively.

Those and further assumptions made by each hypothesis are laid out in Table 1 and Figure 2.

**Figure 1:**
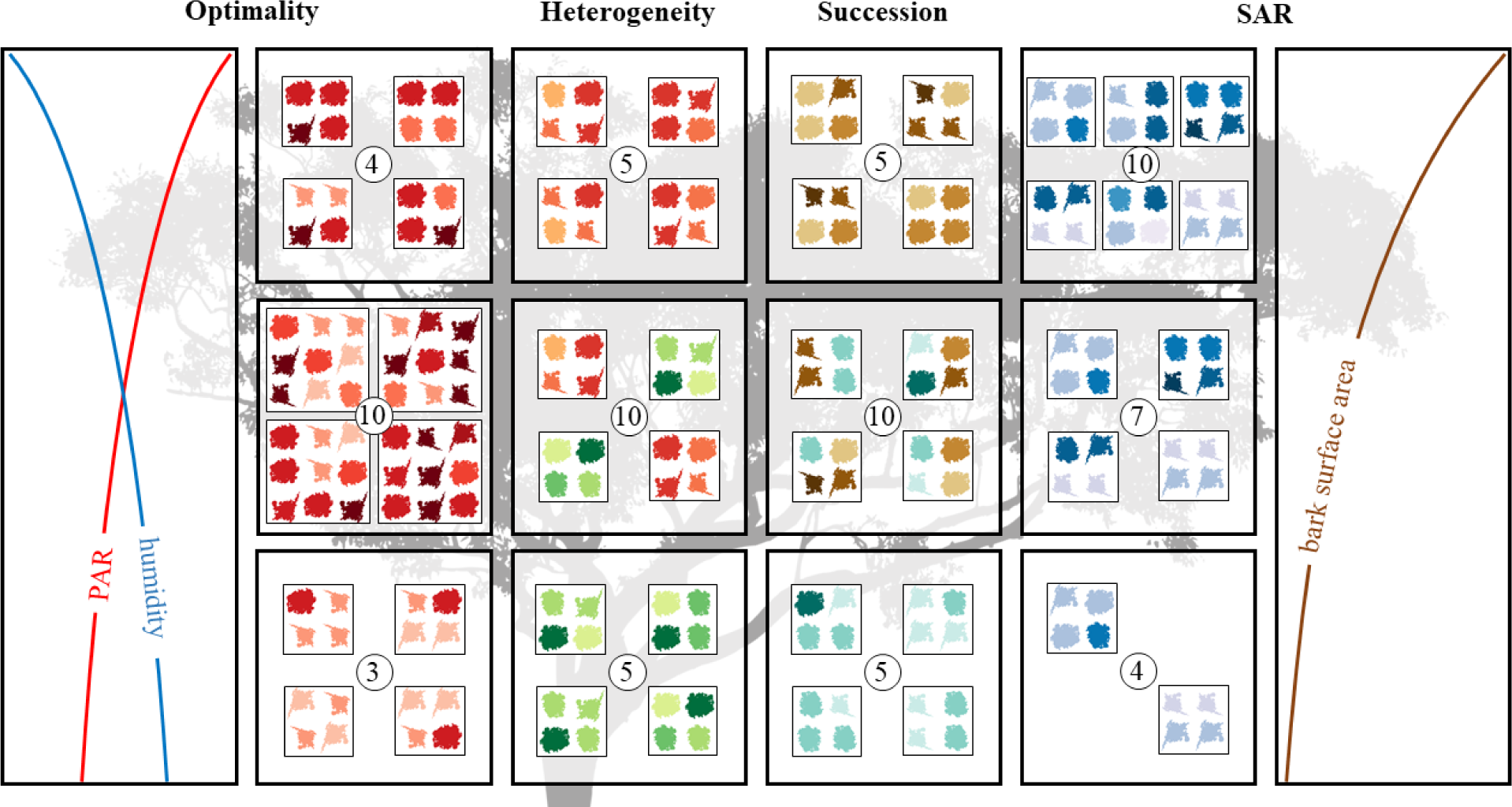
Schematic representation of the interplay between biodiversity shaping mechanisms (columns) and their expected spatial fingerprints on layer-level species richness (γ diversity, represented as encircled number in the center). Each smaller square represents a sampling unit area within a layer (bigger square). Optimality: Concurrence of environmental variables such as available light and humidity leads to favorable growth conditions in a mid-canopy layer. Heterogeneity: Differentially adapted species (indicated by different color gradients) coexist in a more heterogeneous layer in the mid crown. Succession: Early and late successional species (indicated by different color gradients) coincide on branches of median age opposed to young branches in the top crown layer and old branches in the bottom layer. Species Area Relation (SAR): The creation of surface area due to continuous branching along the vertical axis leads to a species-indiscriminate increase in richness.

**Figure 2:**
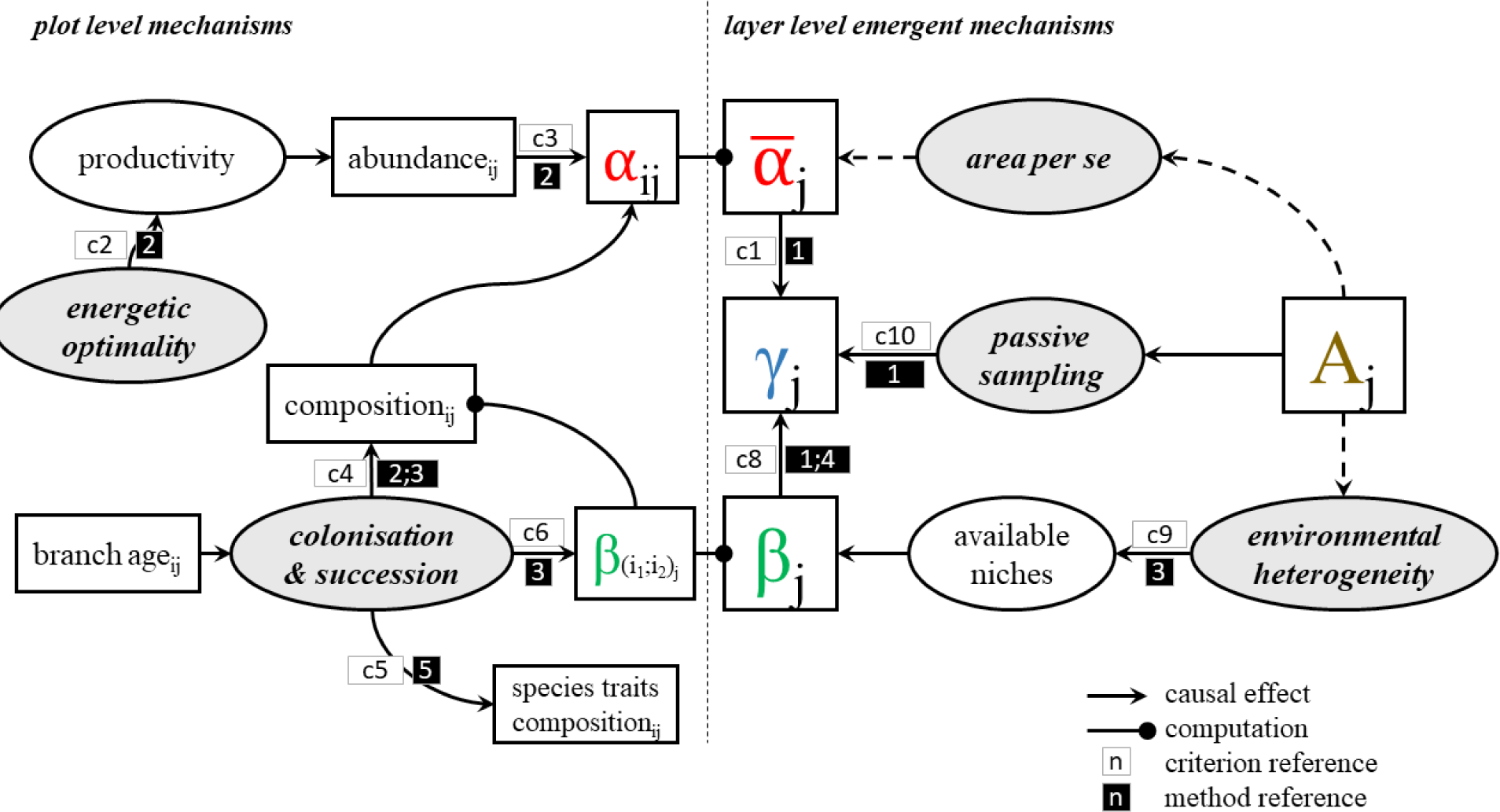
Conceptual scheme of hypothesized mechanisms for the emergence of lichen diversity in the tree crown. Rounded boxes represent the mechanism itself, square boxes measurable variables. Numbers in small white boxes refer to expected conditions in the evidence matrix (Table 1), numbers inside black boxes refer to statistical methods used (detailed in table 2).

**Table 2:** Used statistical methods by reference number (white on black) and criteria from Table 2 to be tested (reference in brackets)

| reference number | model type | Aim |
| --- | --- | --- |
| **1** | multiple linear regression model | test contributions of mean $\alpha$ and $\beta$ diversity and area to $\gamma$ diversity (c1,c8,c10) |
| **2** | piecewise structural equation model | contributions to $\alpha$ diversity (c2,c3,c4) |
| **3** | distance-based redundancy analysis | contributions of environmental variation to beta diversity (c6, c9) |
| **4** | Raup-Crick Null model | delineate $\beta$ diversity characteristics independent of $\alpha$ diversity variation (c7,c8) |
| **5** | variance analysis of trait gradients | test for a successional trait gradient (c5) |

### 2.4 Analysis

Data Analysis was carried out in in R, version 3.5.1 (R Core Team 2018). In this study, species richness on plot level is referred to as α diversity while layer-level species richness is referred to as γ diversity. As real abundances in terms of individual counts are hard to estimate in lichen without resorting to molecular methods (Snäll et al. 2004, Walser et al. 2005, Gjerde et al. 2012), within this study abundances are represented as cover. Within-layer β diversity was calculated on incidence data using the Simpson-based multiple-site dissimilarity (Baselga 2010). A composition index was calculated as first axis of a non-metric multidimensional scaling (nMDS) ordination using the function *metaMDS* implemented in the vegan package (Oksanen et al. 2019). The number of dimensions (k) was deliberately set to one to generate a single compositional gradient (variable ‘com’ in Figure 3). Goodness of fit was evaluated using the *stressplot* function from the same package, in which R^2^ is defined as 1−*S*^2^, with S being the stress value, which is a measure based on the sum of squared residuals of the ordination (Legendre and Legendre 2003 p. 447 ff.). The nMDS ordination was based on a β diversity matrix, which used pairwise between-plot dissimilarities based on the Simpson β diversity index (β_sim_; Koleff, Gaston, & Lennon, 2003).

**Figure 3:**
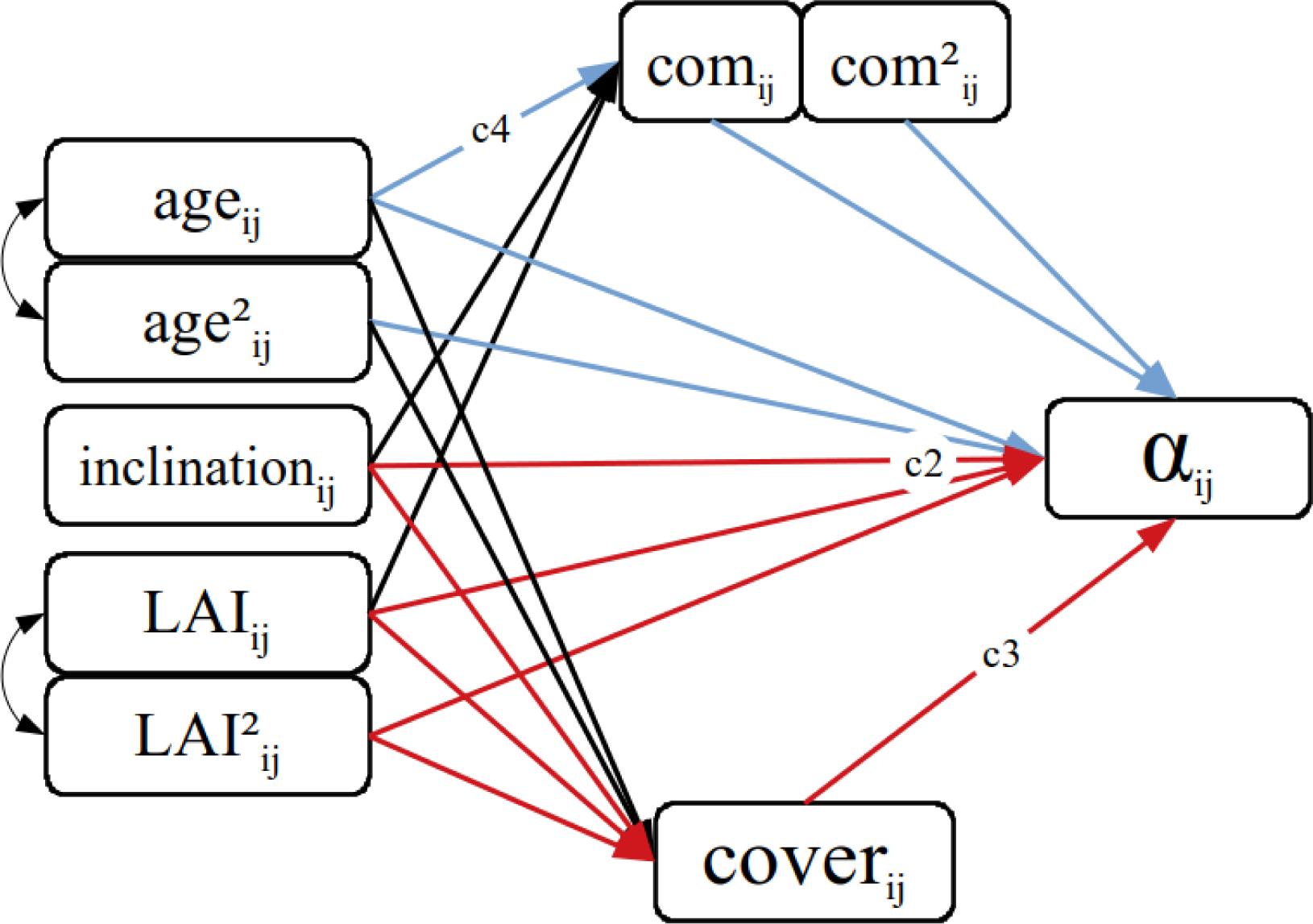
piecewise structural equation model for the plot level. Mechanistic distinctions can be made according to the path followed (The numbers correspond to conditions mentioned in Table 1 and Figure 2). According to the optimality hypothesis, paths in the lower half (2,3; marked red) should be more distinguished as species richness is thought to be driven by energetic optimality and mediated by abundance, represented by leaf area index (LAI) and cover, respectively. If α diversity is mainly determined by successional mechanisms species richness is expected to depend strongly on branch age, mediated by species composition (respective pathways marked blue). For each branch age, LAI and composition index both the linear and quadratic term are included to account for hump shaped patterns. No mechanistic distinction can be stated for either heterogeneity or Species area relation hypothesis as these are assumed to be emergent at the layer level.

#### 2.4.1 Multiple linear regression model

An important distinction between the hypotheses is whether variation in γ diversity and mean α diversity are closely connected (*Optimality, Succession*) or whether patterns in γ diversity are an emergent property at the layer level either driven by variation in β diversity (*Heterogeneity*) or area (*SAR*). To get insight into this question a multiple linear regression model was created using a z-transformed layer-level data set with γ diversity as response variable and mean α diversity, β diversity and available surface area as explanatory variables. Additionally, the interaction terms of area with mean α and β diversity were included. This allowed us to assign a potential area effect to one of its composite mechanisms *heterogeneity* or *area per se* which make different predictions about their scale of impact (Schoereder et al. 2004).

#### 2.4.2 Piecewise Structural Equation Model

To test for the plausibility of aforementioned mechanisms of shaping biodiversity patterns being expressed at the plot level (α diversity) piecewise structural equation modelling (SEM) was applied. This analysis was performed for the pooled phorophyte species dataset and on datasets containing only a single phorophyte species, separately. The SEM model was constructed using the piecewiseSEM package in R (Lefcheck 2016). Numerical data had been either log- or tukey-transformed using the rcompanion package (Mangiafico 2019), if the transformation improved residual normal distribution of the models used within the SEM and subsequently standardized (z-score). Exponents used for transformation are provided in the supplement (S3). Global model fit was evaluated using the Fisher’s C. The Fisher’s C statistic is derived of the combined p-values of each independence claim associated with the hypothesized path diagram of the SEM, known as the *basis set* (Lefcheck 2016). A test of directed separations (Shipley 2013) was performed on the model to test for missing possible paths, which upon inclusion would lead to an improvement of the model. Such improvements of the model were compared against the initial model using the Akaike’s information criterion (AIC) as calculated by Shipley (2013).

The piecewise SEM was constructed with α diversity as main target variable (Figure 3). SEMs feature exogenous variables and endogenous variables, the first of which are variables without incoming causal relations within the model. The latter are defined as being caused by other variables. Endogenous variables may also mediate causal relations to further endogenous variables. Exogenous variables on the plot level include branch age as linear and quadratic term, branch inclination and LAI in both a linear and quadratic term. The inclusion of both linear and quadratic terms for branch age and LAI was based on the expectation of a hump-shaped relation of species richness according to the mid-successional diversity peak and a hump-shaped productivity-diversity relationship, respectively. Originally, bark roughness was included as additional exogenous variable to capture small scale local variation in branch surface environment, but it was dismissed due to its high correlation with branch age (Spearman correlation coefficient = 0.83). All endogenous variables within the plot level are dependent on all the aforementioned exogenous variables. These are the composition index, cover and α diversity. Branch inclination and LAI are assumed to have implications for the energy budget of epiphytic lichen and thus are expected to show significant relations towards α diversity under the *Optimality mechanism* (Figure 2). As under this assumption species richness is regulated via higher population densities a strong link between cover and α diversity can also be expected. Branch age can be expected to be a strong determinant for the sequential establishment of lichen communities along the successional gradient. The composition of these communities in turn is intended to be captured with the composition index. The *Succession mechanism* hypothesis (Figure) predicts an overlap of distinct early and late successional communities at the mid successional peak, which is translated into a link of the community indicator towards α diversity. As the shape of this relationship can expected to be hump-shaped, the quadratic composition index is added as predictor for α diversity as well. Thus, the piecewise SEM is constructed based on three linear models:

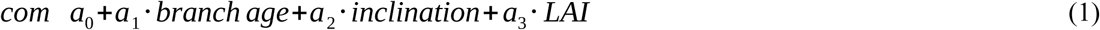

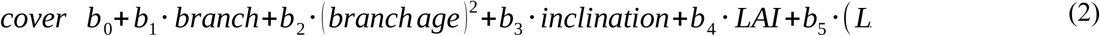

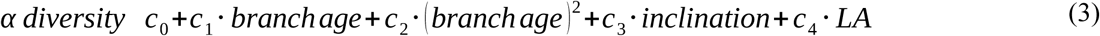

With *com* referring to the composition index and *a, b* and *c* to the estimated slopes of equation 1, 2 and 3, respectively. In order to highlight potential idiosyncrasies of the phorophyte species, this SEM is calculated on both the full data set as well as separately for each species.

#### 2.4.3 Distance-based redundancy analysis

A distance-based redundancy Analysis (dbRDA, Legendre & Anderson, 1999) was applied to the same pairwise β_sim_-matrix, which also served as the basis for the nMDS (see above). Categorical identity properties (phorophyte species, layer) as well as environmental and structural properties (branch age, inclination, bark roughness, LAI, height and distance to the tree center) were used to delineate variation in turnover and community composition both spatially and mechanistically. Variance partitioning (S8) was applied to check for significance and quantify explained variation of these predictors.

#### 2.4.4 Raup-Crick Null model

In studies comparing α and β diversities, both measures are usually expected to be correlated to some degree, as many β metrics can vary simply due to changes in either α or γ diversity (Chase et al. 2011). Null model approaches can be used to disentangle and correct for this correlation. Furthermore, these null models offer insight into the mechanistic signature of β diversities, as the correlation between α diversity and a given β diversity metric offers a null expectation which observed β diversities may fall below or exceed. In our case, β diversities within layer higher than expected may hint towards deterministic species sorting processes between communities as expected with the *Habitat Heterogeneity* mechanism; β diversities within a layer lower than expected may hint towards deterministic sorting processes that operate across layers, for example along a successional gradient. The model used in this study was originally devised by Raup & Crick (1979) and modified by Chase et al. (2011) and has been applied in numerous studies (Muñoz et al. 2004, Zhou et al. 2014, Dini-Andreote et al. 2015). It provides an index ranging from −1 (communities are more similar than expected by random chance) to 1 (communities are more dissimilar than expected by random chance) with a value of 0 corresponding to the Null expectation.

#### 2.4.5 Species trait variance analysis

Successional series of lichens are often reflected by changes in trait composition (Rogers 1990, Lawrey 1991). Early successional species tend to have smaller thalli and reproduce sexually and with smaller diaspores. Late succession species achieve competitiveness through faster growth rates and are defended against herbivores, pathogens and UV radiation by secondary metabolites (Lawrey 1986, Ellis 2012). For all observed lichen species an average value of the composition index could be computed using the presence/absence data for each plot. If the composition index adequately represents a successional gradient, this would yield the species average position on this gradient. These species averages were modelled in a variance analysis against species traits known to reflect a successional trait gradient (Rogers 1990), such as growth form, dispersal strategy and their reaction to chemical identification tests as a proxy for chemical traits (K-,C- and P-test), to check for significant trait gradients.

## 3 Results

A total number of 27 lichen species was recorded (17 ± 2.5 species per individual tree; for a complete list see S4). Out of these, 20 species occurred on both phorophyte species. Four species were exclusive to *F. excelsior* and three species to *Q. robur*.

Species richness patterns were similar between both phorophyte species, differing more between layers than between tree individuals or tree species (Figure 4). In both α and γ diversity a general increase with height could be observed with a sharp decrease in the topmost layer. Trunks were almost devoid of any lichen growth with only 7 species found there and 61.3 % empty plots compared to 3.8 % in all crown layers. Layer 4 was the most species-rich layer in all trees, containing all but 5 of the recorded species. On both plot and layer level, species richness was significantly correlated with lichen cover (Spearman; α: 0.65; γ: 0.68; both p < 0.001). Within-layer β diversity was lowest on the trunks and in layer 5. Values for the inner crown layers were not significantly different from each other (Kruskal-Wallis; p = 0.66; p < 0,001 for all layers). Correlations between the environmental and structural predictors (age, bark roughness, branch inclination and LAI) were generally high and significant, but most pronounced between branch age and bark roughness (Spearman; *Fraxinus*: 0.90, *Quercus*: 0.91; p < 0.001), and branch age and LAI (Spearman; *Fraxinus*: 0.66, *Quercus*: 0.67; p < 0.001).

**Figure 4:**
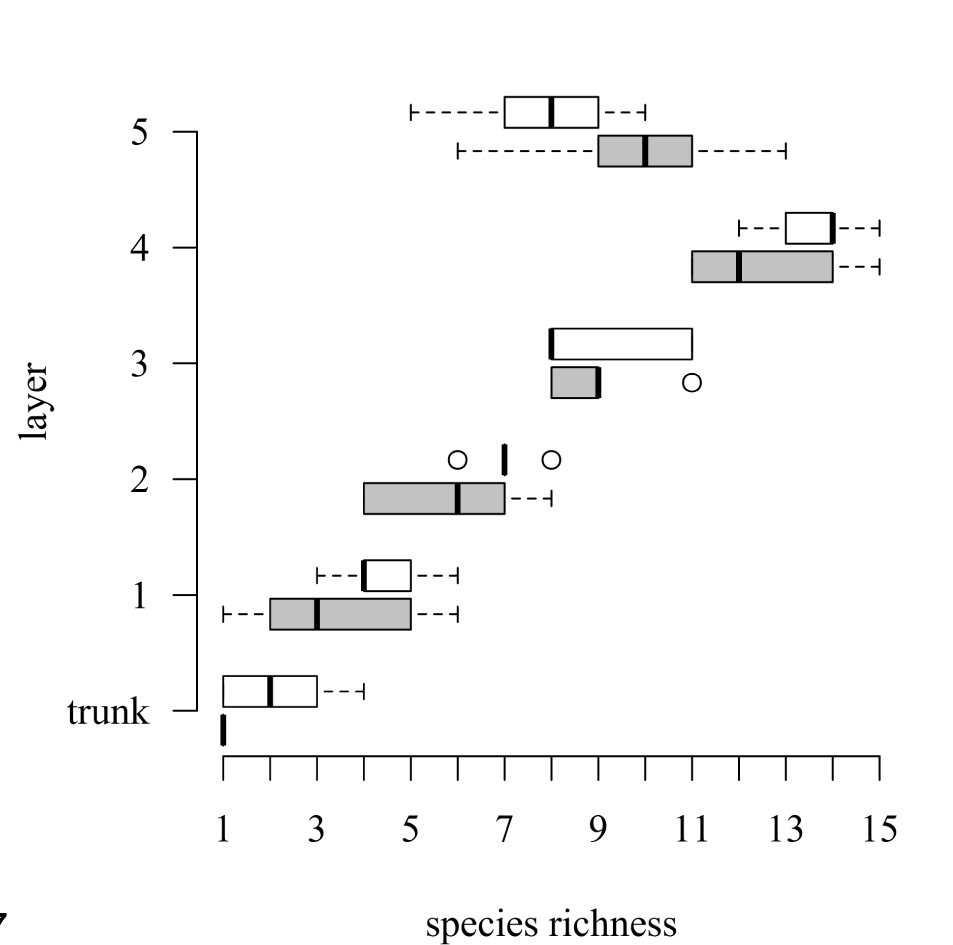
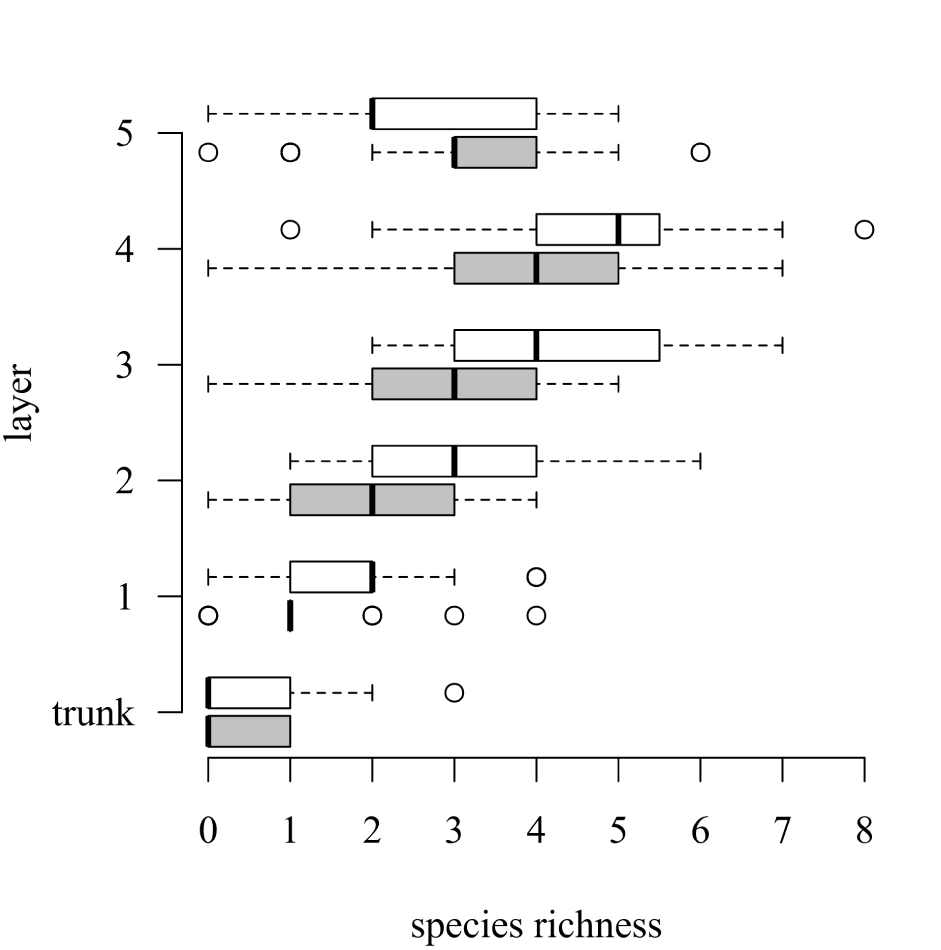
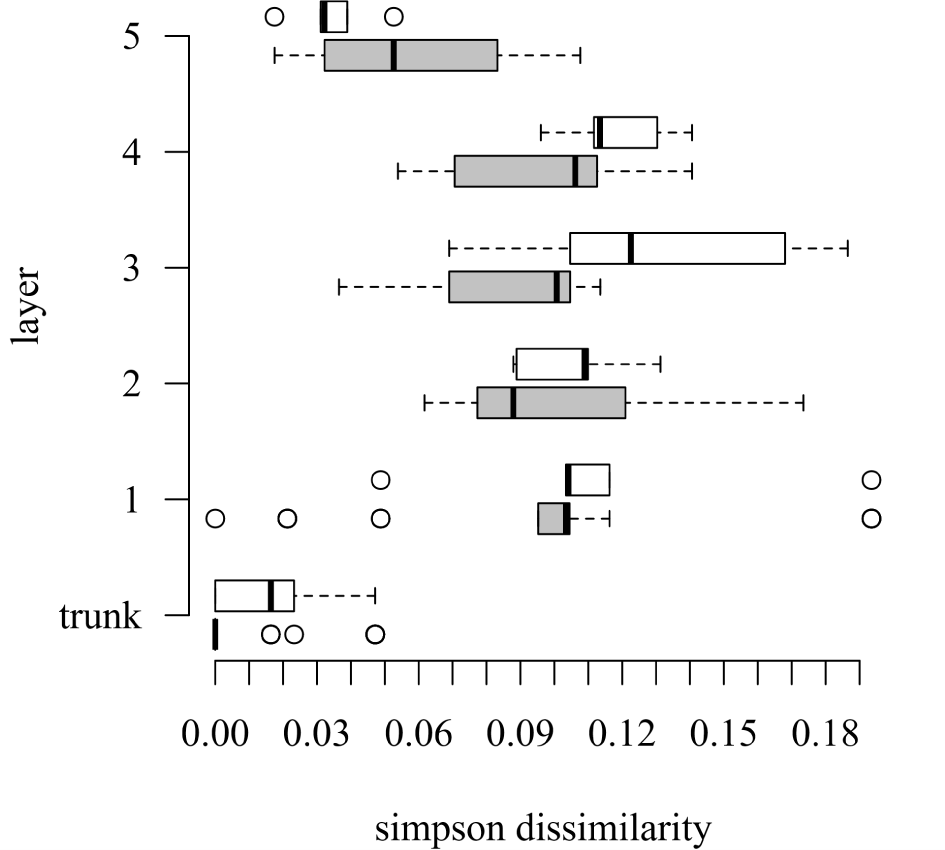
Vertical patterns of lichen diversity in the tree crown. White boxes represent individuals of *Fraxinus excelsior*, grey boxes *Quercus robur*. a) *γ* diversity (layer level) b) α diversity (plot level) c) β diversity (within layer)

### 3.1 Multiple linear regression

In the multiple regression modeling γ diversity as a function of mean α diversity, β diversity and area (including interactions), mean α diversity had the strongest effect (0.87, p < 0.001), with the only other significant effect being the interaction term of α diversity and area (0.31, p = 0.002).

Available surface area (Figure 5) alone only had a small, non-significant effect (0.12, p = 0.10). Overall explained variance was 82.4 %.

**Figure 5:**
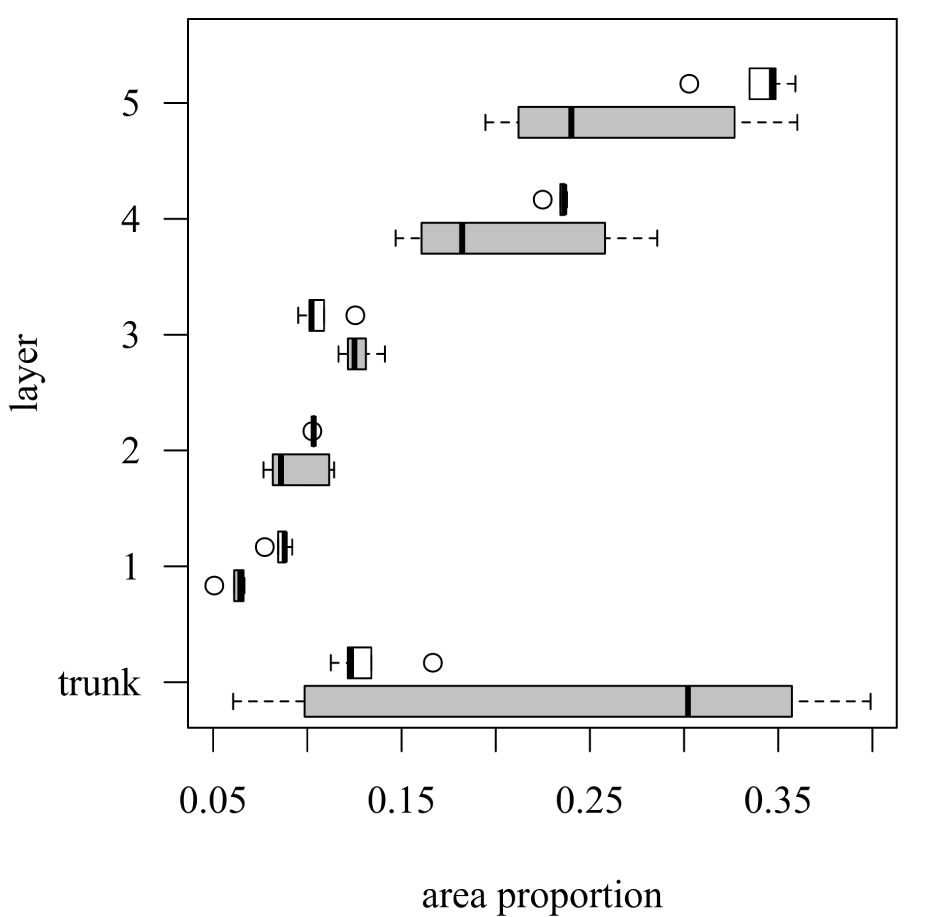
Modelled amount of available surface area within layers in relation to total tree surface. White boxes represent *Fraxinus excelsior*, grey boxes *Quercus robur*. Note the amount of variation in surface area being visibly higher in *Q. robur*.

### 3.2 Piecewise Structural Equation Model

The SEM exploring the determinants of α diversity using the full data set, including both phorophyte species, obtained a non-significant Fisher’s C (8.52, p = 0.384), indicating an appropriate representation of the data. The tree species-specific models performed similarly well (*F. excelsior*: 7.726, p = 0.461; *Q. robur*: 10.784, p = 0.214).

Branch age was the exogenous parameter with the strongest significant effect on all plot-level dependent variables (Figure 6) with concave quadratic relations towards α diversity (−0.20, p < 0.001; linear term: −0.11, p = 0.002) and cover (−0.50, p < 0.001; linear term non-significant) and linear relations towards community composition (−0.67, p < 0.001). Yet, cover constituted the strongest predictor for α diversity (0.63, p < 0.001). LAI had significant effects on α diversity (linear: −0.06, p = 0.033), cover (linear: −0.23, p < 0.001, quadratic: −0.11, p < 0.001) and community composition (linear: −0.08, p = 0.026), while branch inclination only affected community composition significantly (0.09, p = 0.001). The effect of community composition on α diversity was non-significant. Variance explained in endogenous variables was 72 % (α diversity), 40 % (cover) and 52 % (community composition).

**Figure 6:**
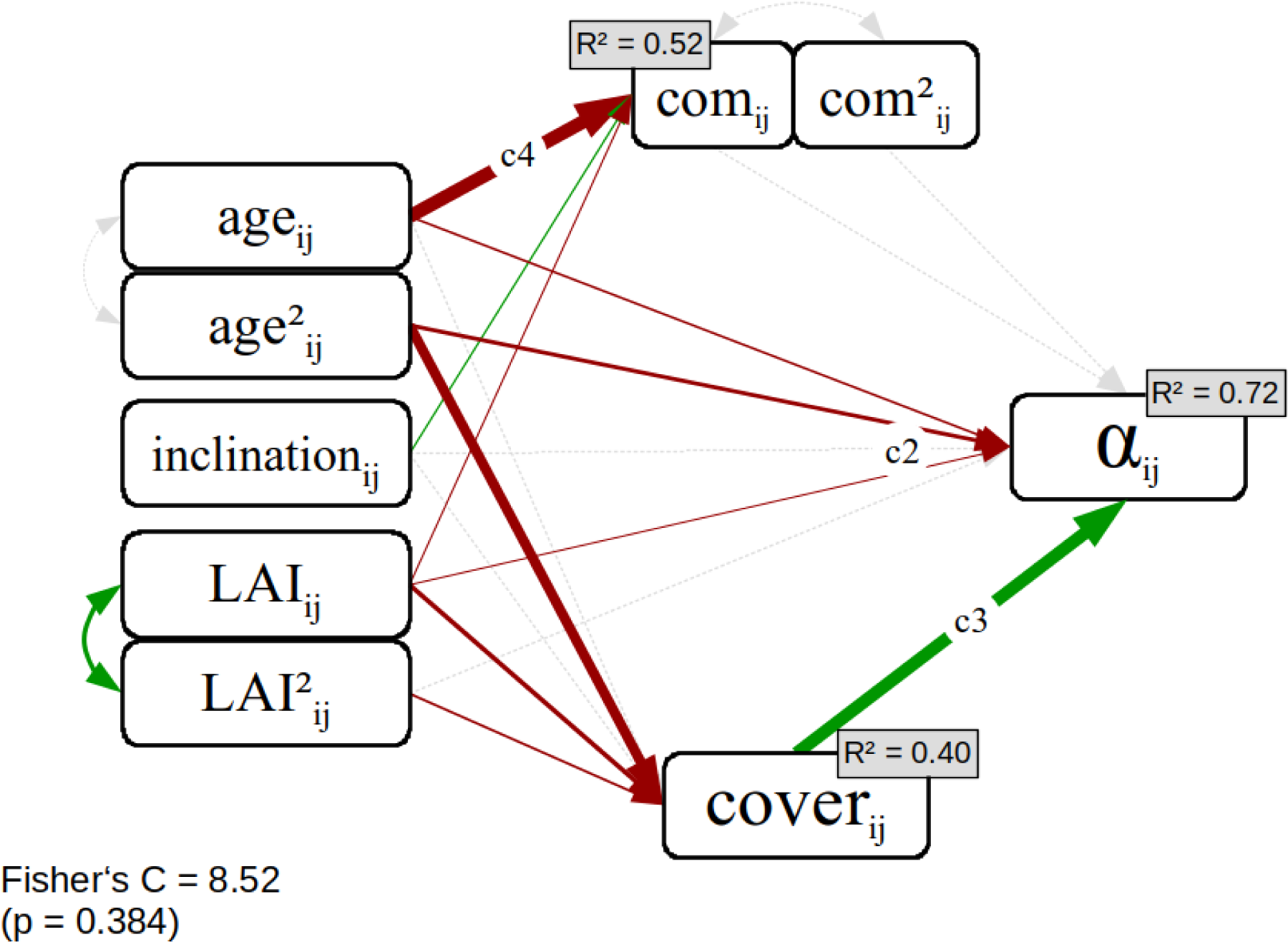
piecewise SEM of plot level pathways using the full dataset. R^2^ in small boxes in top right; arrow width is proportional to parameter estimates, straight lines indicate regressions, double-headed, curved arrows covariance dotted lines p < 0.1, full lines p < 0.05, grey lines p > 0.1 plot marker (j); layer marker (i). See supplement S7 for species specific models.

**Figure 7:**
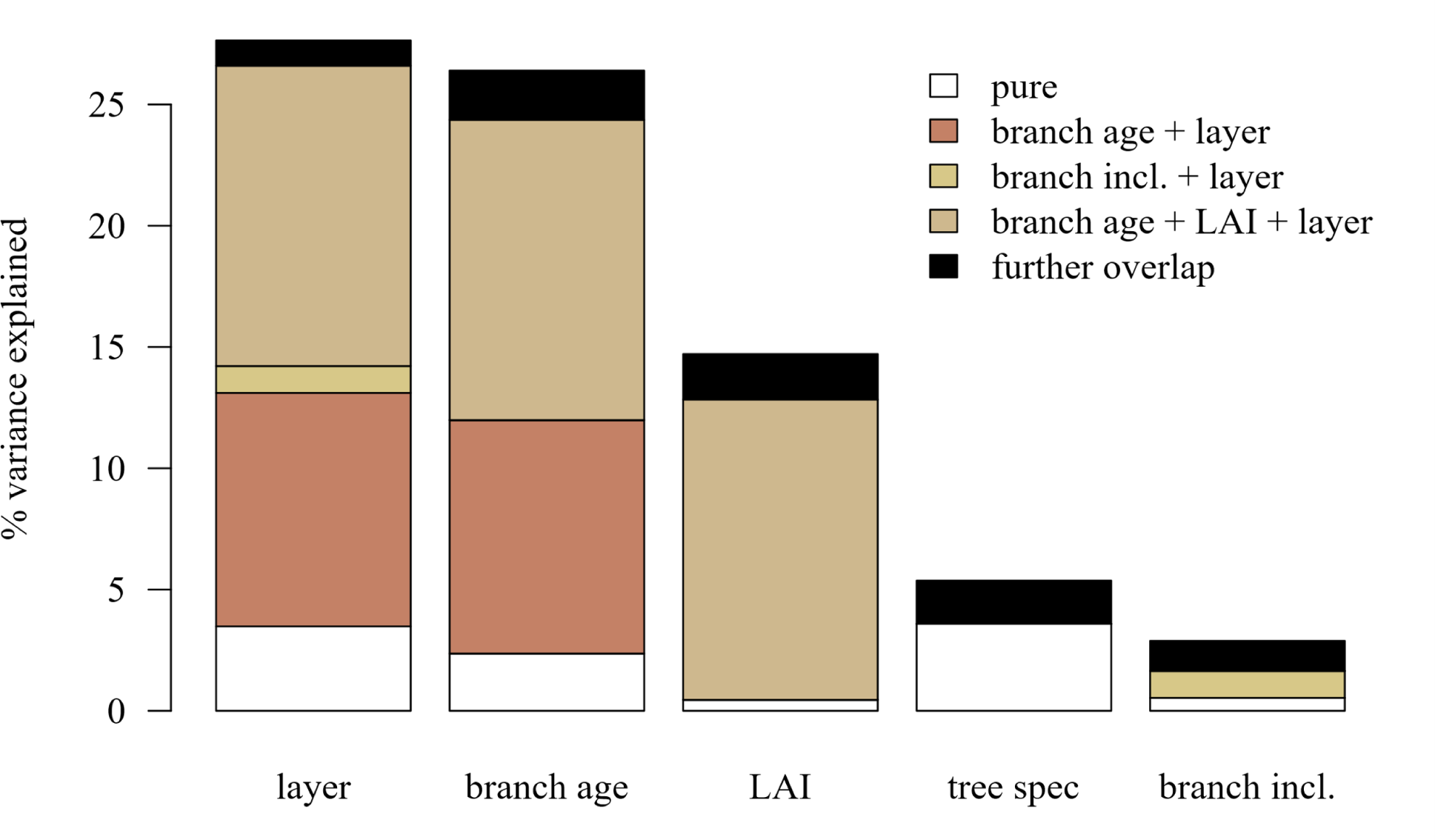
Variation partitioning of β diversity (dbRDA). The bars represent total explained variation by respective variables as the sum of its pure effect and overlap shared with other variables. Blank portions of the bars represent the pure effect; black portions represent the sum of shared overlaps each explaining less than 1% of variance in β diversity. Overall explained variation: 34.9 % (65.1 % unexplained).

The species-specific models (S7) confirmed the main patterns of the full model, although they diverged in a few aspects. While the number of significant paths was slightly reduced in the model for *F. excelsior*, the model for *Q. robur* included one additional significant path compared to the full model, a negative linear effect of branch age on cover. As a consequence, both hump-shaped branch age relations towards α diversity and cover were shifted slightly towards younger branches in *Q. robur* with a maximum of the species richness-branch age relation at 41 ± 6 years compared to 50 ± 3 years in *F. excelsior* (45 ± 6 years in all trees).

### 3.3 Distance-based Redundancy analysis

Pairwise between-plot species turnover (dbRDA, overall 34.9 % explained variance) varied mainly between layers (26.8 % variance explained, 4.0 % purely attributed to layer with no overlap to other effects, see Figure 9), with plots of different layers having a significantly higher β diversity than plots within layers of equal height (Wilcoxon; p < 0.001). All crown environmental variables were significant predictors of pairwise β diversity, such as branch inclination (p = 0.003), LAI (p = 0.010) and especially branch age (p < 0.001), except for bark roughness, which did not explain additional variation in β diversity and was thus removed from the model. They generally shared high overlap with layer. This might imply that these factors are in part responsible for between layer differences. Branch age, for example, explained 25.1 % of variation in pairwise β diversity, 22.7 % of which was shared with layer (which also includes 13.9 % overlap with LAI), while 2.4 % of variation was purely attributed to branch age.

Phorophyte species explained a comparably small amount of variation (5.5 %) but displayed very little overlap with other effects.

### 3.4 Raup-Crick Null Model

Within layers, the number of shared species between plots was significantly higher than null model expectations (t-test, p < 0.001) resulting in mostly negative within-layer means of the Raup-Crick metric (−0.36 ± 0.26, Figure 8). This implies within-layer homogeneity and thus γ diversity control by raised levels of α diversity rather than β diversity. Exceptions could mainly be found on the trunk, where the observed number of shared species mainly matched null model expectations (−0.08 ± 0.10). Deviations from the null model where highest in layer 5 (−0.62 ± 0.12). The increase in deviation from the trunk to the uppermost layers diverged in shape between the tree species with the increase being steeper in *F. excelsior* compared to *Q. robur*.

**Figure 8:**
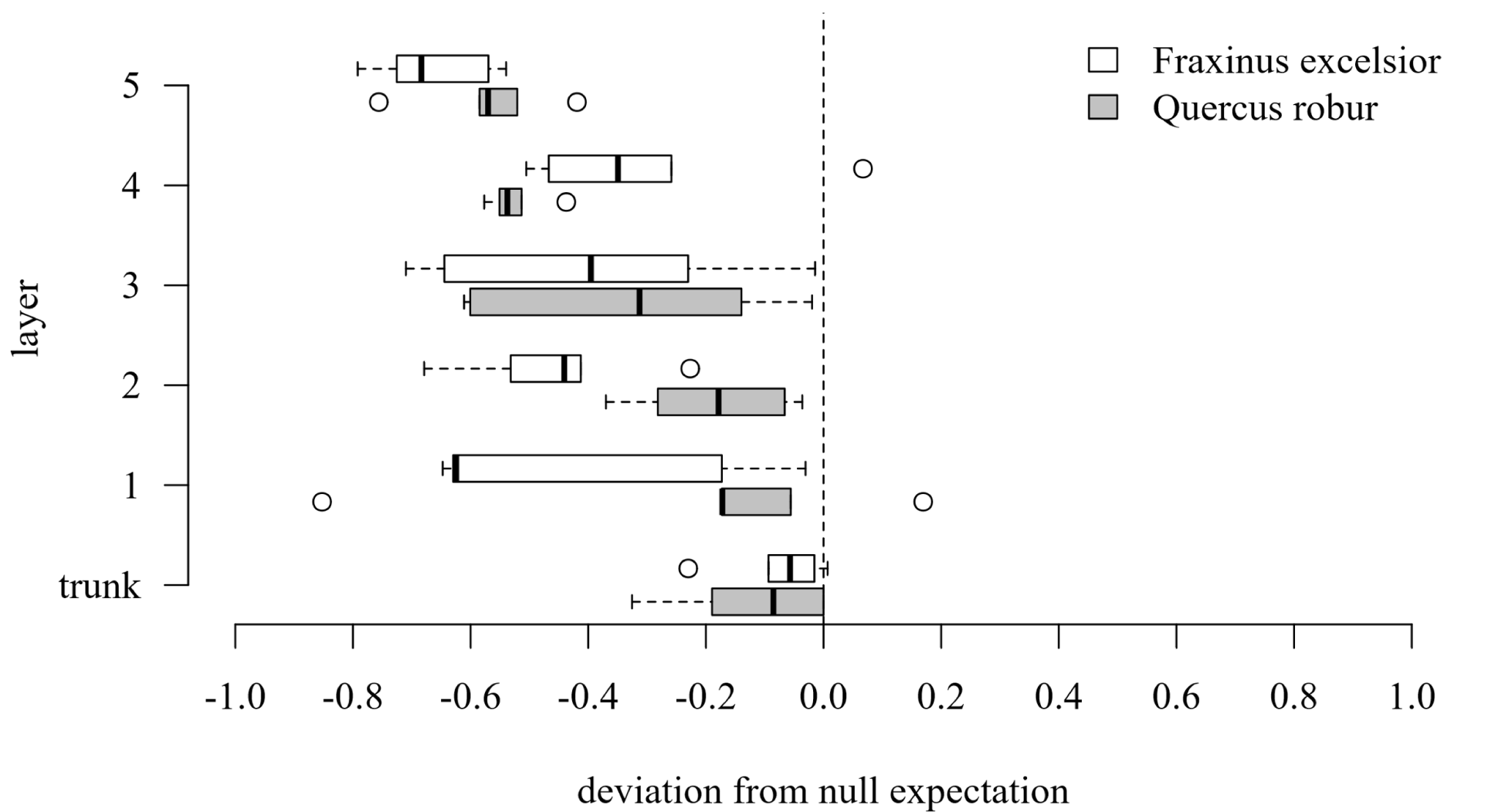
Raup-crick model: mean Raup-Crick metric [-1,1] value by layer and tree species. A value close to 1 represents communities more dissimilar than expected in the null model, indicating community assembly to be highly deterministic (environmental filtering) within layers. Negative values represent communities more similar compared to the null model, indicating assembly processes to be highly deterministic across layers with the communities within the layer being more similar than expected by chance, while values close to zero conform with null expectations and indicate stochastic community assembly.

**Figure 9:**
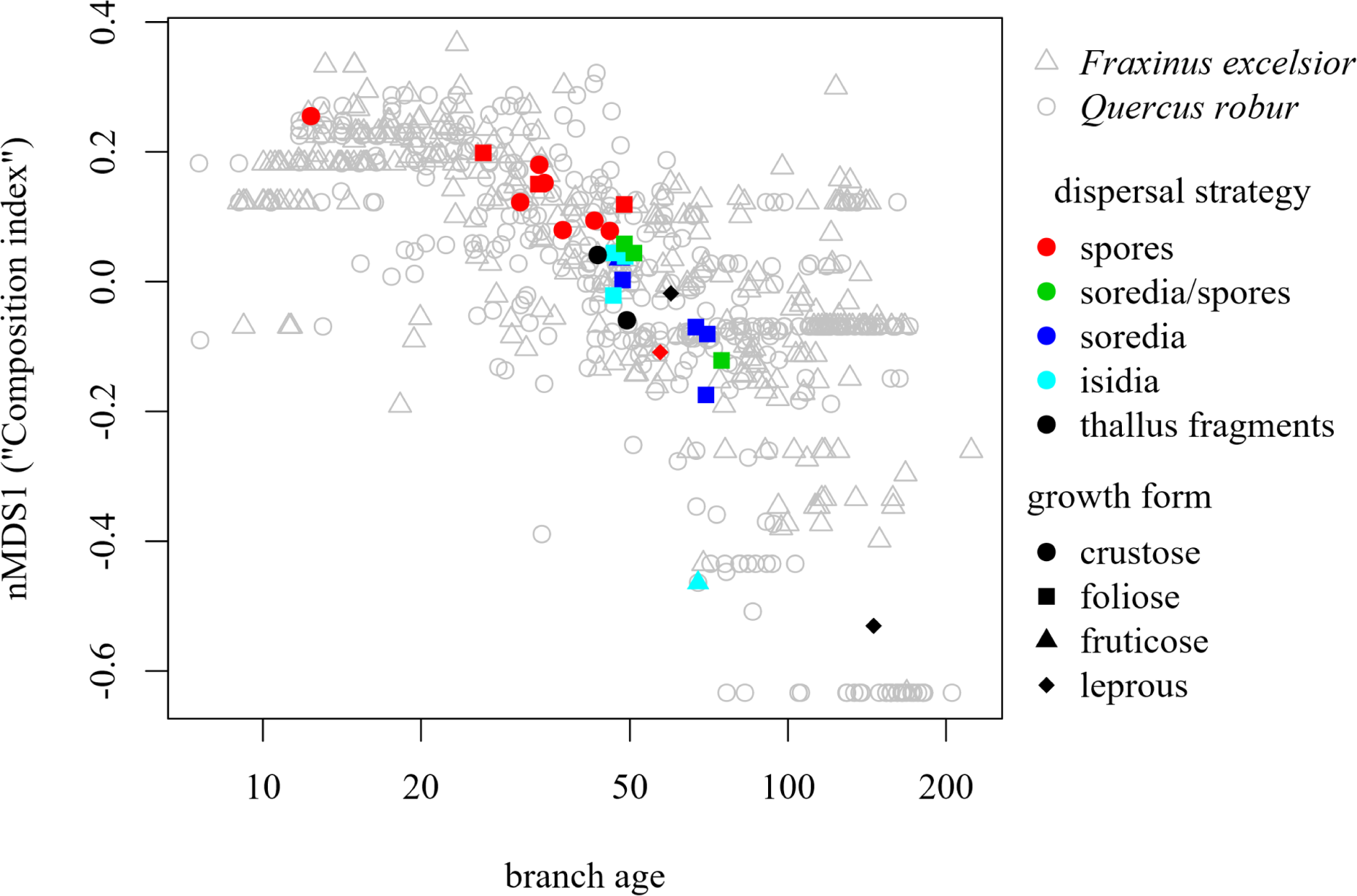
Species trait relations towards branch age. Grey triangles (*Fraxinus excelsior*) and empty circles (*Quercus robur*) represent sampling plots in a 2D composition by branch age ordination. The composition index is calculated as first axis of a non-metric multidimensional scaling (nMDS) based on a pairwise β diversity matrix. Species means for both the nMDS axis as well as branch age determine species position in this 2D ordination, represented by the colored symbols. Noteworthy, a trend for spore-dispersed, crustose and small foliose lichens to be aligned to younger branches, whereas species in which vegetative dispersal is more common are increasingly prevalent with increasing branch age.

### 3.5 Species composition and Species trait variance analysis

Foliose lichens constituted the most common growth form in both species numbers (13) and total cover. Crustose lichens (10) were less abundant. Leprous species were fewer in number of species (3) but had amongst them one of the most abundant species *Candelariella xanthostigma* (Ach.) Lettau. Finally, only three fruticose species could be observed, all of them in small numbers and restricted to only three tree individuals. The nMDS (k=1, n=652) conducted in order to create the composition index produced a stress value of 0.26 (non-metric R^2^ of 0.93). The composition index itself showed negative correlations towards branch age (see fig. 11; Spearman: −0.70; p < 0.001) and LAI (Spearman: −0.51; p < 0.001) but was independent of branch inclination (Spearman: −0.06; p = 0.140). Variations in the averages of the composition index for each species were best explained by growth form (33.3 %, including 16.3 % unshared explained variation; p = 0.002) and presence of Apothecia (30.0 %, including 12.7 % unshared explained variation; p = 0.020). Other traits in regards to dispersal mode or chemistry did not contribute significant explanation of variance in composition (all p > 0.2).

## 4 Discussion

Several lines of evidence have been investigated in order to trace the influence of *Optimality*, *Succession*, *Heterogeneity* and *SAR* as mechanisms shaping patterns in lichen diversity in the tree crown (Table 3). The emergence of the gradients in richness and composition due to these mechanisms are not necessarily mutually exclusive, but may have interactions and added effects, and are discussed in the following subsections. While criteria for the *Optimality* and *Succession mechanisms* were generally met, we could not find any evidence for the *Heterogeneity mechanism*. The *SAR mechanism* was only weakly and indirectly supported: area did not influence γ diversity directly, but had a significant interaction term with α diversity.

**Table 3:**
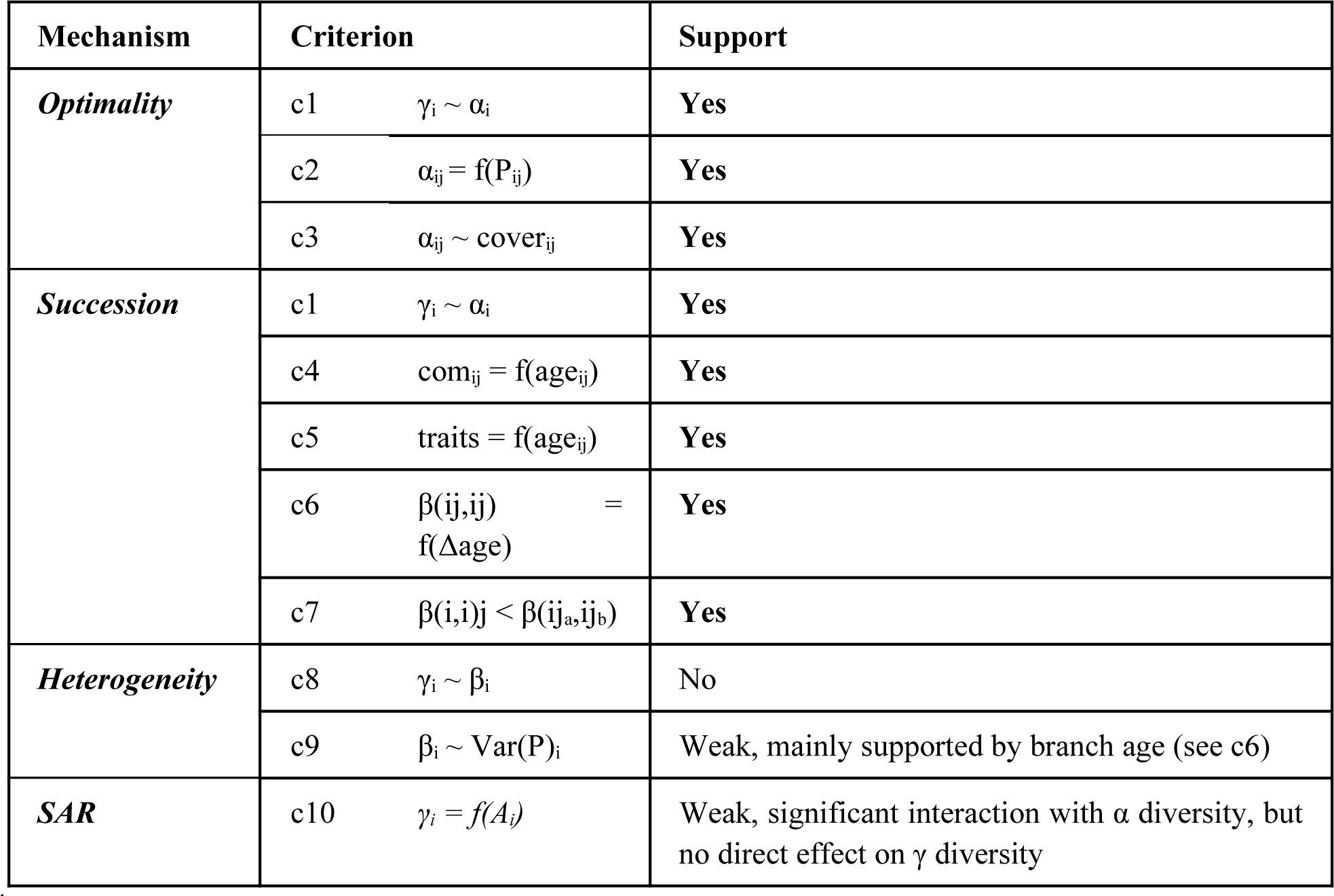
Criteria from the evidence matrix (Table 1) with information about their support by the results. Symbols and abbreviations used are: α, β, γ = α, β or γ diversity; com = community composition; A = branch surface area; P = habitat predictors (e.g. branch inclination, LAI); age = branch age, ‘traits’ is referring to a successional species trait gradient; i = layer marker, j = plot marker (j_a_ and j_b_ mark different layers). The criterion numbers are for reference in Figure 2.

### 4.1 Optimality

Both α and γ diversity exhibited a mid-canopy peak and were significantly correlated (criterion 1, Table 1). In contrast, a distinct peak in β diversity was not observed and β diversity was not correlated with γ diversity. An increase in α diversity was expected to be mediated by an increase in cover (criterion 3, Table 1). This was supported by the SEM, where cover was the strongest predictor of α diversity (Figure 6). Cover itself depended significantly on LAI (criterion 2, Table 1), displaying a hump-shaped relationship. Notably, this includes a decrease of cover on old branches in the dim lower part of the trunk, on which sufficient time for colonization without light limitation could have created a high lichen cover (compare 4.2). Lichens, being photoautotrophic organisms, certainly depend on available light in the canopy. It has been shown to be a significant predictor for species richness of epiphytic lichens (Gustafsson and Eriksson 1995, Fritz et al. 2009, Moning et al. 2009, Normann et al. 2010, Rosabal et al. 2012). However, other microclimatic factors like water limitation can be of overriding importance (Sillett and Antoine 2004, Rambo 2010). While we did not measure humidity, we expected that water limitation is highest in the top crown due to drying winds and higher irradiation leading to suboptimal growing conditions thus creating the observed hump-shape. The strong signal of *Optimality* emphasizes the importance of growing seasons conditions for lichen diversity patterns, because during the leaf-off period neither light nor humidity gradients are particularly pronounced.

### 4.2 Succession

In the piecewise SEM, branch age was not only an important determinant of species richness (Figure 10) but also of species composition (criterion 4, Table 1). It even outperformed LAI as predictor of both cover and richness. After accounting for height layer and phorophyte species, pairwise differences in composition between plots (dbRDA, see 3.3) also were best explained by branch age (criterion 6, Table 1). Moreover, species composition was expected to be more similar within layers than between layers (criterion 7, Table 1), which was supported by both the dbRDA variance partitioning and the Raup-Crick null model deviations. In the first case, the majority of variation was attributed to height layers. In the second case, excluding the trunk, within-layer deviations of the Raup-Crick null model were mostly negative, indicating that plots within layers were more similar than expected by chance.

**Figure 10:**
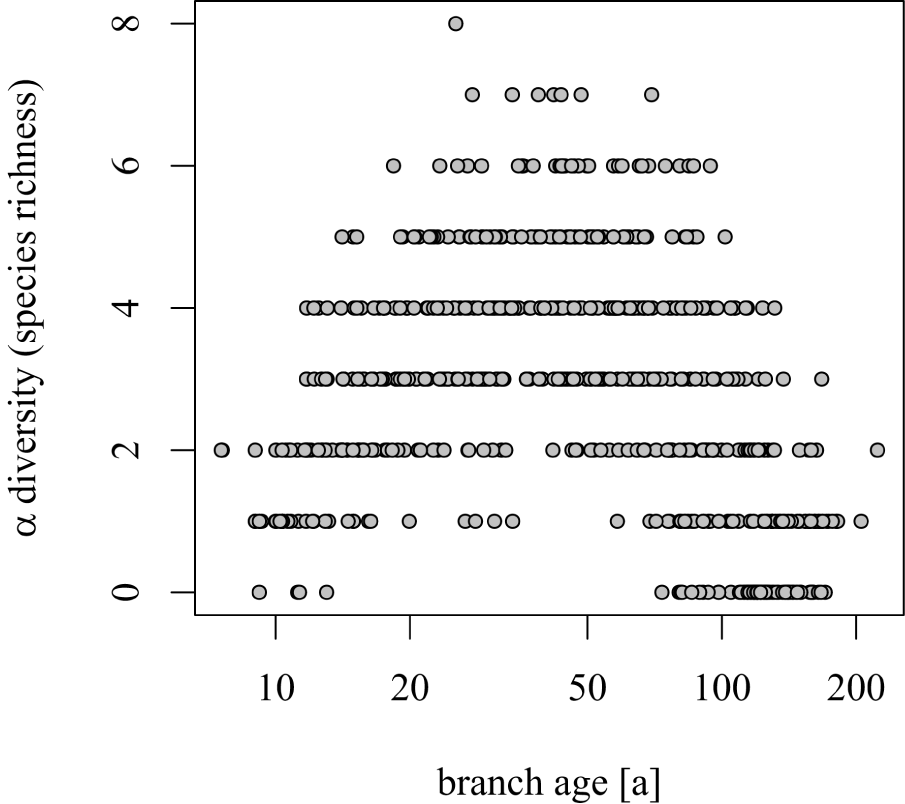
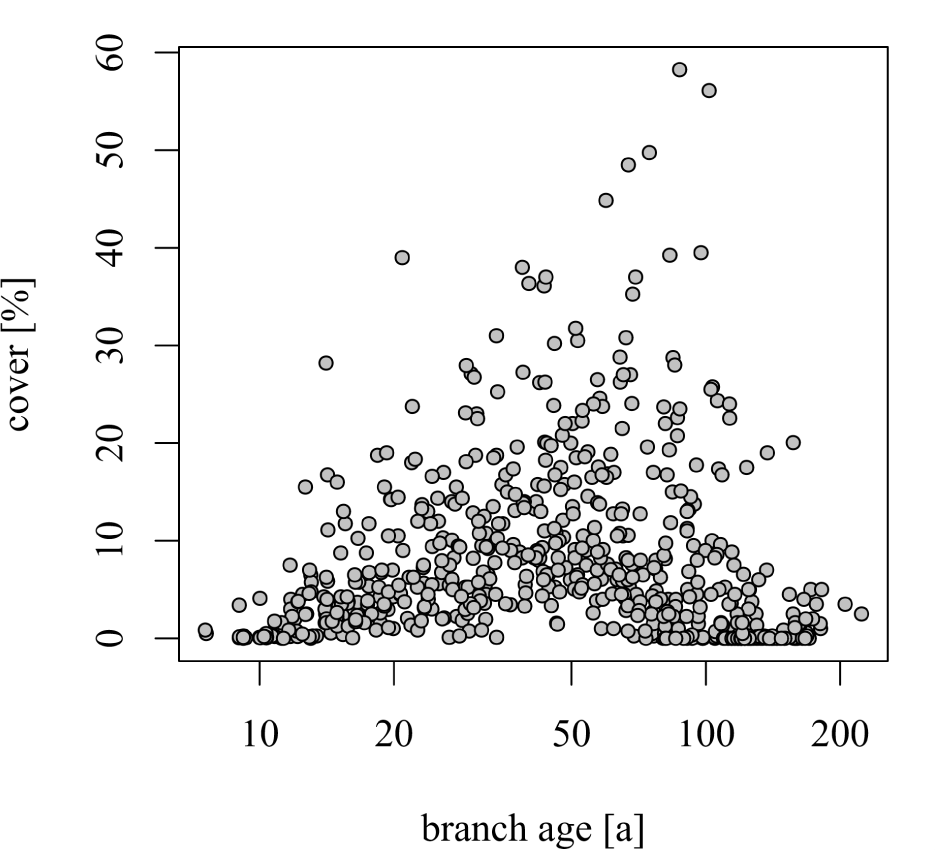
Both the α diversity and cover - branch age relationship shows a distinct hump shape

Evidence for succession further requires branch age to be connected to a successional sequence of epiphytic lichen (Degelius 1964, Hilmo 1994, Wirth et al. 1999), expressing a systematic trait gradient (criterion 5, Table 1). Most common pioneers comprise of physically smaller crustose or foliose lichen with lower investments in longevity. These typically settle on the first uneven surfaces like bud scars and joints (Degelius 1964, Rogers 1990, Ellis 2012). This could be observed in this study and growth form was the trait that best explained the compositional gradient. Common species found in the top canopy layer included small, intricate foliose species (i.e. genus *Physcia* and *Xanthoria*) and crustose species like *Amandinea punctata* (Hoffm.) Coppins & Scheid. Smooth branch surfaces were mainly colonized by small rosettes of crustose species (e.g. genus *Lecanora* and *Caloplaca*). Another typical trait for pioneers is the ability to efficiently disperse propagules in order to colonize new sites (Rogers 1990, Armstrong and Welch 2007). The presence of apothecia did also explain the compositional gradient, with species developing apothecia being more prevalent on younger branches. Apothecia produce sexual spores that are much smaller and hence are dispersed more readily than asexual modes of dispersal (soredia, isidia and thallus fragments). Bigger, more robust foliose species, fruticose and leprous species with predominantly asexual modes of dispersal appeared further down on relatively older branches and the trunk.

One caveat remained: The hump-shaped relation between branch age and α diversity (Figure 11) might suggest a mid-succession peak at a branch age of about 45 years. However, the decline in species richness on the peak’s descent is assumed to result from competitive exclusions (Purschke et al. 2013), also described as “quorum effect” (Jenkins 2006). Evidence for competitive interactions, such as overgrowth between lichens could not be observed on the lower parts of the crown and the trunk. On the contrary, lichen thalli became more sparsely distributed and the cover decreased (Figure 10b) making competitive exclusion less probable. This is in line with previous studies on epiphytic lichens which concluded that competitive exclusions rarely occurs once thalli are established (Lawrey 1991, Snäll et al. 2003, Pentecost 2014).

In conclusion, the *Succession* mechanism aptly describes the increase in species richness as environmental filters get alleviated and species accumulate on the surfaces of growing branches that are gradually overtopped and sheltered as the tree gains height. Past the species richness maximum, however, a different mechanism than competitive exclusion has to account for the decrease in cover and species richness on the oldest parts of the tree. As possible alternative mechanisms competition by bryophytes, shedding of bark surface (Cáceres et al. 2007) or grazing pressure by gastropods (Asplund et al. 2010) have been suggested. However, the coinciding decrease in environmental *Optimality* and of lichen cover on older branches in lower canopy layers despite the long time for colonization, hinting towards less favorable conditions further below (see 4.1) might be the strongest candidate.

### 4.3 Heterogeneity

Heterogeneity is amongst the first and the most commonly documented drivers of diversity (Stein et al. 2014). The horizontal variation of environmental variables has been highlighted as distinct ecological feature of tree crowns (Parker and Brown 2000) and as driver of arboreal diversity patterns. This has been exemplified for tropical vascular epiphytes (Woods et al. 2015). It is thus surprising that in our study heterogeneity showed little to no influence in shaping the diversity gradient within the trees.

Firstly, β diversity was not related to γ diversity (criterion 8, Table 1) and secondly, deviations from the Raup-Crick null model did not indicate that within-layer differences were higher than expected by chance. Both would have been prerequisites for the *Heterogeneity mechanism*. Although variation in pairwise β diversity was reasonably well explained in the dbRDA, little variation was attributed to within-layer variation of environmental variables (criterion 9, Table 1). Out of 34.9 % explained variation, 26.2 % could at least partially ascribed to height layers, 4.4 % to phorophyte species, leaving only about 4 % remaining to be explained.

### 4.4 Species Area Relationship

In the multiple linear regression analysis, only mean α diversity and its interaction with area had a significant effect on γ diversity. Available surface area was not significant as a main effect (criterion 10, Table 1). Thus, area does not have an effect at the layer level, but it amplifies the effect of α diversity in layers with a high amount of surface area. This may suggest that area does have an influence on small scale processes of species richness regulation in line with the *area per se* – hypothesis (Connor and McCoy 2000, Schoereder et al. 2004). However, layers in the upper crown not only possess greater branch surface area; they also are less shaded and span a greater extent of projected area. As a result, they are able to collect more photosynthetically active radiation. The interaction effect of surface area may thus also contain elements of the *Optimality mechanism*.

### 4.5 Conclusion and implications for lichen biodiversity in canopies

The results of this study suggest an intricate interplay between the mechanisms of environmental *Optimality* and *Succession* in controlling vertical diversity patterns of epiphytic lichen in the tree crown. This includes environmental limitations in the harshly exposed young branches and dim understory (*Optimality*), as well as the strong vertical species turnover associated with successional dynamics in the upper height layers. On older branches in height layer four both mechanisms jointly create a sheltered, yet light habitat where a majority of species of the whole successional gradient co-occur to create the observed peak in species richness. The strong increase in surface area with height resulting from the fractal branching process did not translate into a monotonously increasing diversity signal. While we found horizontal heterogeneity within height layers to have little effect, vertical heterogeneity in environmental conditions and successional time appeared to be strong driver of lichen diversity and composition in our study (compare Figure 7) and in the literature (Bates 1992, McCune et al. 2000). These gradients may be steeper than elevational gradients by an order of magnitude (Nakamura et al. 2017) and thus have the potential to strongly contribute to the creation of the biodiversity hotspots that are forest canopies (Ozanne et al. 2003, Nakamura et al. 2017).

### 4.6 Outlook

It comes as little surprise that the effects of mechanisms such as *Succession* and *Optimality* were hard to disentangle. Not only because they are found to be mechanistically intertwined, but so are the predictors used to describe energetic as well as age relations. LAI and branch age were significantly correlated as the architecture of the tree itself is responsible in shaping the vertical gradients found within the canopy (McCune et al. 2000). Such correlations put limits on the insights that can be gained in observational studies. Experiments which separate both effects, for instance by distributing standardized logs as lichen habitat along the vertical and horizontal gradient (Antoine and McCune 2006), offer a way to separate and quantify the relative importance of mechanisms like *Succession*, *Optimality* as well as surface area relationships and dispersal. Effects of dispersal limitations were not included in the original hypotheses as lichen are considered to have a generally high dispersal ability at the canopy scale we considered (Muñoz et al. 2004, Lenoir et al. 2012). However, evidence for small scale dispersal limitations does exists, in particular in dependence of propagule size (Sillett et al. 2000, Löbel et al. 2006). We propose to install air traps collecting diaspores and use the knowledge of lichen distribution to infer dispersal kernels depending on diaspore size. Finally, lichens are a ubiquitous element of biodiversity in temperate tree crowns but by far only one of many. To date, only limited literature exists on multi-taxa approaches on multi-trophic dynamics (Shorrocks et al. 1991, Nadkarni 1994, Lamit et al. 2015, Asplund et al. 2016). It certainly would be of interest to investigate how the patterns of lichen diversity found within this study propagate through the complex ecosystem that is the forest canopy including diversity patterns of lichen-associated organisms as well as processes such as nutrient cycling. Facilities such as the Canopy Crane in Leipzig are promising tools to tackle this challenge.

## Supporting information

compiled supplemental material, 8 tables and 1 R script

## Acknowledgments

We would like to thank Antonia Ludwig, Joy Opitz and Cordelia Weis for their help with fieldwork, Dr. Peter Otto for providing access to the *herbarium universitatis lipsiensis* and sharing his expertise on determining lichens. We further thank Prof. Dr. Volkmar Wirth for helpful comments and his pioneering contributions to the field of lichen taxonomy and ecology without which this study would not have been possible and Christian Vonderach for the provision of the dataset of *random branch sampling*, which allowed us to quantify gradients in available branch area, and finally the German Centre for Integrative Biodiversity Research (iDiv) for supporting the Leipzig Canopy Crane Facility.

