## Supplementary material for "Tree crowns as meeting points of diversity generating mechanisms – a test with epiphytic lichens in a temperate forest": compiled supplemental material, 8 tables and 1 R script

### Supplement 1: Dirichlet Regression Model

To account for surface area available in each height layer of the sampled trees, we modelled bark surface area by means of a Dirichlet Regression. This regression was based on a dataset of 385 broadleaf trees which was obtained utilizing the *random branch sampling* method (Riedel and Kändler 2017). The *F. excelsior* model used diameter at breast height (DBH) as predictor for a second-order polynomial model, the *Q. robur* model used a logarithmic model with DBH as predictor and the ratio crown base height / total tree height as additional predictor. The parameter estimates and significance levels of the regression coefficients for these regressions are detailed in table S1.

**Table S1:** Summary outputs of the Dirichlet-regressions modelling the bark surface area A in layer i as proportion of total bark surface area in the whole tree as a function of diameter at breast height and the crown base height/total height ratio (p). The type of model is depicted below its corresponding tree species.
Number of observations: 385; Significance levels: < 0.001 ‘***’, 0.001 ‘**’, 0.01 ‘*’, 0.05 ‘.’.

***Fraxinus excelsior***
$A_i = \beta_0 + \beta_1 \cdot DBH + \beta_2 \cdot DBH^2$

| **Layer** | | trunk | 1 | 2 | 3 | 4 | 5 |
|---|---|---|---|---|---|---|---|
| **estimate** | **Intercept** | 0.567 | -0.743 | -1.158 | -1.127 | -1.076 | -0.645 |
| | ***DBH²*** | $-6.91 \cdot 10^{-6}$ | $-3.46 \cdot 10^{-6}$ | $-5.98 \cdot 10^{-6}$ | $-8.48 \cdot 10^{-6}$ | $-6.84 \cdot 10^{-6}$ | $-5.71 \cdot 10^{-6}$ |
| | ***DBH*** | $5.44 \cdot 10^{-3}$ | $3.89 \cdot 10^{-3}$ | $6.63 \cdot 10^{-3}$ | $8.58 \cdot 10^{-3}$ | $8.24 \cdot 10^{-3}$ | $7.28 \cdot 10^{-3}$ |
| **p-value** | **Intercept** | 0.003 ** | 0.001 ** | <0.001*** | <0.001*** | <0.001*** | 0.002 ** |
| | ***DBH²*** | <0.001*** | 0.024 * | <0.001*** | <0.001*** | <0.001*** | <0.001*** |

| | | | | | | | |
|---|---|---|---|---|---|---|---|
| | ***DBH*** | <0.001*** | 0.002 ** | <0.001*** | <0.001*** | <0.001*** | <0.001*** |

***Quercus robur***
$A_i = \beta_0 + \beta_1 \cdot \log(DBH) + \beta_2 \cdot p$

| layer | | Trunk | 1 | 2 | 3 | 4 | 5 |
|---|---|---|---|---|---|---|---|
| **estimate** | **Intercept** | 0.010 | -1.756 | -2.672 | -2.164 | -3.565 | -3.367 |
| | **log(*DBH*)** | 0.049 | 0.366 | 0.630 | 0.567 | 0.886 | 0.887 |
| | ***p*** | 3.209 | -0.155 | -1.110 | -0.609 | -1.172 | -1.040 |
| **p-value** | **Intercept** | 0.983 | 0.002** | <0.001*** | <0.001*** | <0.001*** | <0.001*** |
| | **log(*DBH*)** | 0.540 | <0.001*** | <0.001*** | <0.001*** | <0.001*** | <0.001*** |
| | ***p*** | <0.001*** | 0.602 | <0.001*** | 0.017* | <0.001*** | <0.001*** |

### Supplement 2: Branch Age Model

Branch age was modelled by means of a multiple linear regression using branch diameter and height on the tree (when standing), their interaction and tree species as predictors. This regression was based on a dataset of 829 branch cross-cuts taken from wind felled *Quercus robur* and *Fraxinus excelsior* individuals in the forest surrounding the Leipzig Canopy Crane Facility. As the dataset was heavily skewed towards small diameters, weights were introduced using kernel density estimation (function kde2d in R). The parameter estimates and significance levels of the regression coefficients for these regressions are detailed in table S2.

**Table S2:** branch age model; The branch age model was calculated as a multiple linear regression model following the formula: $\sqrt{branch\,age} \quad \log(diameter)+species+(diameter:height)$.
$R^2$: 0.80 (adjusted $R^2$: 0.79); Residual standard error: 6.394 on 712 degrees of freedom; F-statistic: 922.8 on 3 and 712 DF, p-value: < 2.2e-16; Significance levels: < 0.001 '***', 0.001 '**', 0.01 '*', 0.05 '.'.

| | **Estimate** | **p-value** |
|---|---|---|
| **Intercept** | -1.376 | < 0.001*** |
| **log(diameter)** | 1.803 | < 0.001*** |
| **species (Q. robur)** | 5.762e-01 | < 0.001*** |
| **diameter:height** | 2.400e-04 | < 0.001*** |

### Supplement 3: Transformation of variables (piecewise Structural Equation Model)

The variables used in the piecewise Structural Equation Model (pSEM, section 2.4.2) were transformed if the procedure improved residual normal distribution of the constituent linear models. This transformation was either achieved by exponentiation or natural logarithm. In case of negative values in a variable x, a term δ, corresponding to the minimal value in x, was added to x prior to exponentiation to prevent loss of data points. The exponents used for transformation (λ) and added terms δ are detailed in table S3.

**Table S3:** Data transformation as used in the pSEM followed the pattern $(x+\delta)^{\lambda}$(except for uses of logarithm).

| **x** | **Λ** | **δ** |
|---|---|---|
| **α diversity** | 0.925 | 0 |
| **branch age** | 0.200 | 0 |
| **branch inclination** | natural logarithm | 2 |
| **composition index** | 2.050 | 0.63 |

### Supplement 4: List of lichen species found at the Leipzig Canopy Crane Facillity

**Table S4:** Species found within this study, sorted by total frequency in all trees. *Physcia adscendens* and *P. tenella* were grouped into *Physcia sp.* when soredia were not yet established. Additionally, frequencies in *F. excelsior* and *Q. robur* trees are given in column 5 and 6, respectively. Two species are listed that have been identified in the studied trees but fell outside the sampling plots.

| **Species** | **Family** | **Growth form** | **frequency** | ***F. excelsior*** | ***Q. robur*** |
|---|---|---|---|---|---|
| ***Physcia sp.*** | Physciaceae | Foliose | 695 | 305 | 390 |
| ***- identified as P. adscendens*** **(Fr.) H. Olivier** | Physciaceae | Foliose | 233 | 67 | 166 |
| ***- identified as P. tenella*** **(Scop.) DC.** | Physciaceae | Foliose | 182 | 90 | 92 |
| ***Candelariella xanthostigma*** **(Ach.) Lettau** | Candelariaceae | Leprose | 259 | 115 | 144 |
| ***Xanthoria parietina*** **(L.) Th.Fr.** | Teloschistaceae | Foliose | 239 | 141 | 98 |
| ***Phaeophyscia orbicularis*** **(Neck.) Moberg** | Physciaceae | Foliose | 142 | 79 | 63 |
| ***Amandinea punctata*** **(Hoffm.) Coppins & Scheid.** | Caliciaceae | Crustose | 140 | 81 | 59 |
| ***Massjukiella polycarpa*** **(Hoffm.) Rieber** | Teloschistaceae | Foliose | 125 | 49 | 76 |
| ***Parmelia sulcata*** **Taylor** | Parmeliaceae | Foliose | 70 | 22 | 48 |
| ***Candelaria concolor*** **(Dicks.) Stein** | Candelariaceae | Foliose | 59 | 32 | 27 |
| ***Lepraria incana*** **(L.) Ach.** | Stereocaulaceae | Leprose | 40 | 5 | 35 |
| ***Physcia stellaris*** **(L.) Nyl.** | Physciaceae | Foliose | 27 | 21 | 6 |
| ***Melanohalea exasperatula*** **(Nyl.) O. Blanco, A. Crespo, Divakar, Essl., D. Hawksw. & Lumbsch** | Parmeliaceae | Foliose | 22 | 13 | 9 |
| ***Hypogymnia physodes*** **(L.) Nyl.** | Parmeliaceae | Foliose | 15 | 9 | 6 |
| ***Melanelixia fuliginosa*** **(Fr. ex Duby) O.Blanco et al.** ***subsp. glabratula*** **(Lamy) J.R.Laundon** | Parmeliaceae | Foliose | 13 | 3 | 10 |
| ***Punctelia subrudecta*** **(Nyl.) Krog** | Parmeliaceae | Foliose | 12 | 5 | 7 |

| | | | | | |
|---|---|---|---|---|---|
| ***Lecanora hagenii subsp. persimilis*** **Th. Fr.** | Lecanoraceae | Crustose | 11 | 11 | 0 |
| ***Lecidella elaeochroma*** **(Ach.) M. Choisy** | Lecanoraceae | Crustose | 7 | 4 | 3 |
| ***Lecanora symmicta*** **(Ach.) Ach.** | Lecanoraceae | Crustose | 7 | 3 | 4 |
| ***Lecanora pulicaris*** **(Pers.) Ach.** | Lecanoraceae | Crustose | 6 | 2 | 4 |
| ***unidentified sterile grey crustose lichen*** | | Crustose | 5 | 1 | 4 |
| ***Evernia prunastri*** **var.** ***herinii*** **(P.A. Duvign.) D. Hawksw.** | Parmeliaceae | Fruticose | 4 | 3 | 1 |
| ***Flavoparmelia caperata*** **(L.) Hale** | Parmeliaceae | Foliose | 4 | 3 | 1 |
| ***Caloplaca cerinella*** **(Nyl.) Flagey** | Teloschistaceae | Crustose | 2 | 2 | 0 |
| ***Lecanora carpinea*** **(L.) Vain.** | Lecanoraceae | Crustose | 2 | 2 | 0 |
| ***Lecanora conizaeoides*** **Nyl. ex Cromb.** | Lecanoraceae | Leprose | 1 | 0 | 1 |
| ***Usnea hirta*** **(L.) Weber ex F. H. Wigg.** | Parmeliaceae | Fruticose | 1 | 0 | 1 |
| ***Cladonia sp. (cf. fimbriata*** **(L.) Fr.*)*** | Cladoniaceae | Fruticose | (not found within plots) | | |
| ***Pseudevernia furfuracea*** **(L.) Zopf** | Parmeliaceae | Fruticose | (not found within plots) | | |

### Supplement 5: Results from the multiple linear regression model (γ diversity)

**Table S5:** Summary of the results from the multiple linear regression model explaining layer-level γ diversity on a scaled layer-level dataset. Predictors were: mean plot-level richness per layer (mean α), Simpson-based multiple-site dissimilarity ($\beta_{sim}$) per layer, available surface area and the interaction terms between area and mean α diversity and β diversity respectively. $R^2$: 0.83 (adjusted $R^2$: 0.82); Residual standard error: 0.4201 on 54 degrees of freedom; F-statistic: 56.06 on 5 and 54 DF, p-value: < 0.001; Significance levels: < 0.001 ‘***’, 0.001 ‘**’, 0.01 ‘*’, 0.05 ‘.’.

| | **Estimate** | **p-value** |
|---|---|---|
| **(intercept)** | -0.085 | 0.190 |
| **mean α** | 0.875 | < 0.001 *** |
| **$\beta_{sim}$** | -0.062 | 0.478 |
| **area** | 0.118 | 0.099 |
| **mean α:area** | 0.313 | 0.002 ** |
| **$\beta_{sim}$:area** | -0.046 | 0.594 |

### Supplement 6: Supporting information on Species Area Relationship in height layers

To account for potential biases due to the varying sample sizes within layer, species richness was extrapolated for a common sample size of 50 replicates ($\gamma50$). Species accumulation curves were interpolated using the method devised by (Coleman 1981) and used for extrapolation via linearized logarithmic function models. To test for species-area relationships the semi-logarithmic model $\gamma = z \cdot \log(A) + c$ was used. $\gamma$ diversity was tested against absolute available surface area. Additionally, the relation was tested for the $\gamma50$ estimate. The results of these models are depicted in Table S6a and Figure S6b.

Replacing $\gamma$ diversity with its $\gamma50$ estimate diminished the effect and the slope. A reduction of the slope in the Species Area Relation when replacing recorded $\gamma$ diversity with $\gamma50$ may hint towards a sampling effect, as recorded $\gamma$ diversity reflects both sampling design and ecological processes, while the effect of sampling design has been removed from the estimate (Cam et al. 2002). This sampling effect may constitute either an artifact of sampling design or effect of passive sampling. Larger areas can be thought of as larger samples of the same environment, thus making passive sampling an important Null hypothesis (Connor and McCoy 2000, Cam et al. 2002). Using species richness estimates assumes incomplete species detection. However, due to their lower architectural complexity in the trunk and lower crown layers species are not only less likely to avoid detection, but 50 plots are close to exceeding the actual available space. In this way even complete sampling would have produced a similar observation. On the other hand, reducing the sample size to a common lower size would certainly have increased the number of undetected species.

**Table S6a:** Summary of the results from the linear regression model testing a layer-level semi-logarithmic Species Area Relation ($\gamma = z \cdot \log(A) + c$). The relationship was tested on datasets only containing information from *Fraxinus excelsior* and *Quercus robur* individuals as well a full data set containing information from both phorophyte species.

Additionally, each relationship was tested for the γ50 estimate. The first row details the amount of explained variance, the second row the significance level (< 0.001 '***', 0.001 '**', 0.01 '*', 0.05 '.') and the third row the slope of the relationship.

| | Whole data set | | *Fraxinus excelsior* | | *Quercus robur* | |
|---|---|---|---|---|---|---|
| | γ diversity | γ50 estimate | γ diversity | γ50 estimate | γ diversity | γ50 estimate |
| $R^2$ | 0.117 | 0.009 | 0.205 | 0.029 | 0.088 | 0.012 |
| p value | 0.007 * | 0.463 | 0.012 . | 0.368 | 0.111 | 0.912 |
| z (slope) | 1.534 | 0.697 | 3.424 | 1.830 | 1.367 | 0.123 |

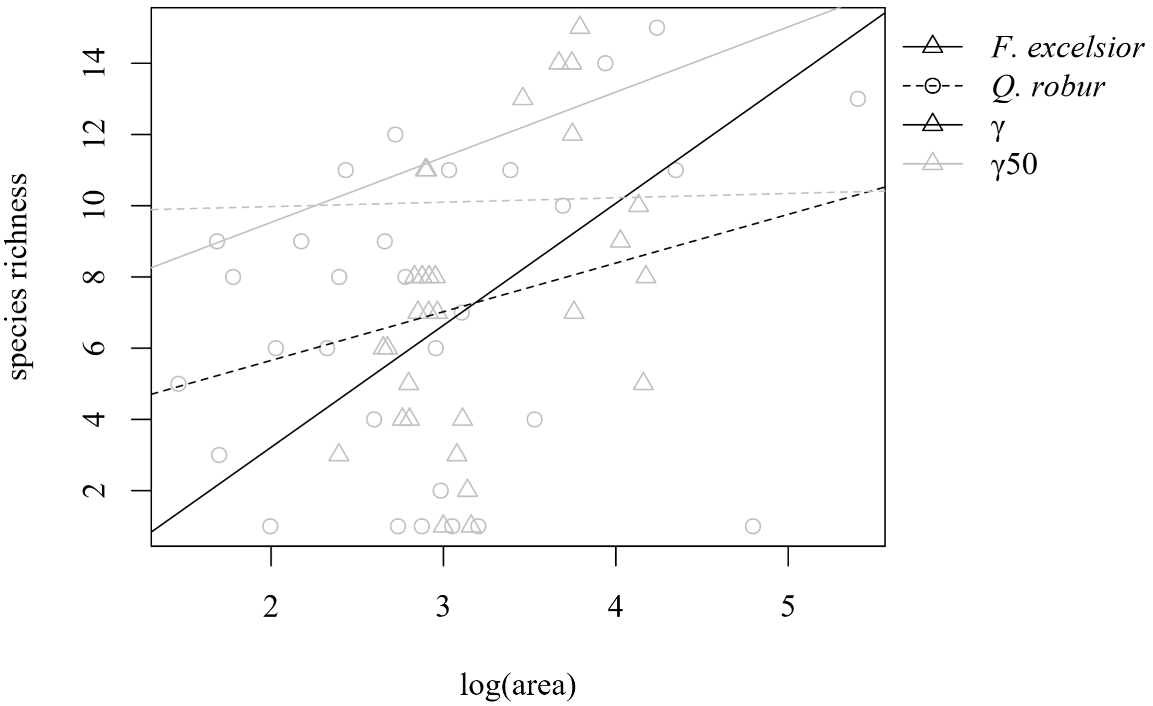


**Figure 6b:** Species Area relationships within the tree crown. The black lines represent semi-logarithmic regressions of observed γ diversities ($\gamma = c \cdot \log(A)^z$), grey lines represent semi-logarithmic regressions of the γ50 estimate ($r^2$ in brackets). Triangles and continuous lines represent Fraxinus excelsior, circles and dashed lines represent Quercus robur.

### Supplement 7: Detailed results of the piecewise Structural Equation Models

Additional to performing the piecewise Equation Model (pSEM) analysis on a plot-level data set containing both phorophyte species (see section 2.4.2), it was also performed on data sets containing only a single species. The results of these analyzes can be found in table S7.

**Table S7:** Summary outputs of the piecewise Structural Equation Models (pSEM) performed on the full plot-level data set and on data sets from only a single phorophyte species. Overall model fit is represented by 'Fisher's C' in the first row. The table is further broken down into its constituent models, showing explained variance ($R^2$) and parameter estimates and p values for each independent variable (significance levels: < 0.001 '***', 0.001 '**', 0.01 '*', 0.05 '.'). 'α' refers to plot level species richness, 'com' stands for composition index, 'age' for branch age, 'LAI' for Leaf Area Index, representing the light conditions in each plot and 'incl.' describes the branch inclination.
In the case of the *Q. robur* pSEM the test of directed separations revealed a significant path (p = 0.048) from '$LAI^2$' towards 'com'. The inclusion of the path mentioned decreased both the AIC of the model by 3.22 and the Fisher's C (5.567, p=0.473). The estimate for this additional path was relatively low (0.055) and improved explained variance of 'com' only by 1 %.

| | | **Full data set** | ***Fraxinus excelsior*** | ***Quercus robur*** |
|---|---|---|---|---|
| **Fisher's C** | | 8.524 | 7.726 | 10.784 |
| **p value** | | 0.384 | 0.461 | 0.214 |
| **$\alpha \sim$ cover + com + age + $age^2$ + LAI + $LAI^2$ + incl.** | | $R^2 = 0.72$ | $R^2 = 0.73$ | $R^2 = 0.71$ |
| **cover** | **estimate** | 0.622 | 0.538 | 0.656 |
| | **p value** | < 0.001 *** | < 0.001 *** | < 0.001 *** |
| **com** | **estimate** | 0.054 | 0.083 | 0.003 |
| | **p value** | 0.081 . | 0.092 . | 0.943 |
| **$com^2$** | **estimate** | -0.014 | -0.014 | -0.042 |
| | **p value** | 0.493 | 0.711 | 0.102 |
| **age** | **estimate** | -0.111 | -0.095 | -0.128 |
| | **p value** | 0.002 ** | 0.080 . | 0.011 * |
| **$age^2$** | **estimate** | -0.202 | -0.303 | -0.161 |
| | **p value** | < 0.001 *** | < 0.001 *** | < 0.001 *** |
| **LAI** | **estimate** | -0.064 | -0.072 | -0.081 |
| | **p value** | 0.033 * | 0.128 | 0.041 * |
| **$LAI^2$** | **estimate** | -0.025 | -0.035 | -0.018 |
| | **p value** | 0.140 | 0.183 | 0.413 |
| **incl.** | **estimate** | -0.040 | -0.075 | -0.009 |
| | **p value** | 0.085 . | 0.055 | 0.759 |

| | | | | |
|---|---|---|---|---|
| **cover ~ age + age$^2$ + LAI + LAI$^2$ + incl.** | | $R^2 = 0.40$ | $R^2 = 0.53$ | $R^2 = 0.40$ |
| **age** | **estimate** | -0.036 | 0.080 | -0.201 |
| | **p value** | 0.310 | 0.209 | < 0.001 *** |
| **age$^2$** | **estimate** | -0.499 | -0.678 | -0.3578 |
| | **p value** | < 0.001 *** | < 0.001 *** | < 0.001 *** |
| **LAI** | **estimate** | -0.233 | -0.256 | -0.183 |
| | **p value** | < 0.001 *** | < 0.001 *** | < 0.001 *** |
| **LAI$^2$** | **estimate** | -0.118 | -0.133 | -0.109 |
| | **p value** | < 0.001 *** | < 0.001 *** | < 0.001 *** |
| **incl.** | **estimate** | -0.028 | -0.096 | 0.003 |
| | **p value** | 0.401 | 0.076 . | 0.931 |
| **com ~ age + LAI + incl.** | | $R^2 = 0.52$ | $R^2 = 0.48$ | $R^2 = 0.56$ |
| **age** | **estimate** | -0.669 | -0.603 | -0.734 |
| | **p value** | < 0.001 *** | < 0.001 *** | < 0.001 *** |
| **LAI** | **estimate** | -0.082 | -0.006 | -0.149 |
| | **p value** | 0.026 * | 0.915 | 0.003 ** |
| **incl.** | **estimate** | 0.092 | 0.016 | 0.149 |
| | **p value** | 0.001 ** | 0.717 | < 0.001 *** |
| **covariances** | | | | |
| **LAI ~~ LAI$^2$** | **estimate** | 0.227 | 0.272 | 0.195 |
| | **p value** | < 0.001 *** | < 0.001 *** | < 0.001 *** |
| **age ~~ age$^2$** | **estimate** | -0.015 | -0.196 | 0.176 |
| | **p value** | 0.706 | 0.001 ** | < 0.001 *** |
| **com ~~ com$^2$** | **estimate** | -0.094 | 0.126 | -0.220 |
| | **p value** | 0.010 ** | 0.018 * | < 0.001 *** |

### Supplement 8: Variance Partitioning R script

In order to determine which parameters best explain the pairwise differences in species composition between plots a distance-based redundancy analysis was conducted (section 2.4.3). As many of the parameters are under the influence of the same strong vertical gradients and thus correlated, additional variance partitioning was performed using the code in Table S8.

**Table S8** R script for Variance/Variation partitioning of lm()-type linear regression models and anovas. Output is a table giving the amount of variance explained by each model parameter individually and as overlap with other parameters.

```
#Y=Target variable OR distance matrix (e.g. output of dbRDA variance
partitioning) OR lm output
#p=predictor variables (character vector, size n, NOT needed if Y is
lm output)
#data=Data.frame containing predictor variables (not needed if Y is
lm output)
#scale (logical): should the data be z-transformed?
#minus.null (logical): Convert negative variance partitions to zero?

VarPart<-function(Y,p,data,adjust=TRUE,scale=FALSE,minus.null=FALSE){
  #should the data be z-transformed?
  if(scale==TRUE){
    data<-data.frame(scale(data[,c(Y,p)]))
  }

  if(missing(data)){
    ifelse(class(Y)=="lm",data<-Y[["model"]],warning("source data not
defined"))
  }

  if(missing(p)){
    ifelse(class(Y)=="lm",{
      p<-colnames(data)[-1]
    response<-strsplit(as.character(Y[["call"]][2])," ")[[1]][1]
      },warning("predictor variables not defined"))
  }

  # combination of all explanatory variables
  combo<-lapply(1:length(p),function(m){combn(p,m)})

  #calculation of Rsquared and adjusted Rsquared for any combination
```

```
of explanatory variables
  r2.res<-sapply(combo,function(a){
    apply(a,2,function(b){

      #formula for the individual models
      form<-as.formula(paste(ifelse(
        class(Y)=="lm",response,ifelse(
          is.dist(Y),"Y",Y)),"~",paste(b,collapse="+")))

      #type of response data (dbRDA vs linear model)
      ifelse(is.dist(Y),
             {mod.res<-dbrda(form,data,na.action = na.omit)
             },{mod.res<-lm(form,data,na.action = na.omit)
             })
      unlist(c(paste(b,collapse="+"),RsquareAdj(mod.res)))
    })})
  r2.res<-do.call(cbind,r2.res)
  r2.res<-
data.frame(predictor=r2.res[1,],expl.Var=as.numeric(r2.res[2,]),adjus
t=as.numeric(r2.res[3,]))
  ifelse(adjust==TRUE,pre<-r2.res$adjust,pre<-r2.res$expl.Var)
  r2.ttl<-pre[nrow(r2.res)]
  prd.all<-strsplit(as.character(r2.res$predictor),"\\+")
  prd.n<-prd.all[[nrow(r2.res)]]

  expl<-vector(mode="numeric",length=nrow(r2.res))
  nexpl<-vector(mode="numeric",length=nrow(r2.res))
  varP<-vector(mode="numeric",length=nrow(r2.res))

  #Calculate the pure partitions and overlaps
  for(o in r2.res$predictor){
    prd.o<-strsplit(as.character(o),"\\+")[[1]]
    combo.o<-unlist(sapply(1:length(prd.o),function(m){
      apply(combn(prd.o,m),2,paste,collapse = "+")
    }))
    ttl.p<-prd.n[!prd.n%in%prd.o]
    expl[r2.res$predictor==o]<-sum(varP[r2.res$predictor%in
%combo.o&r2.res$predictor!=as.character(o)])
    nexpl[r2.res$predictor==o]<-
sum(pre[r2.res$predictor==paste(ttl.p,collapse = "+")])
    varP<-r2.ttl-expl-nexpl
  }

  #If there are negative partitions show them as zeroes?
  if(minus.null==TRUE){varP[varP<0]<-0}
  r2.res$varP<-varP

  #with negative partitions, the total sum of partitions might not
equal the total explained variance of the full model
  if(sum(r2.res[,3][r2.res[,3]>0])!=r2.ttl){warning("total explained
variance != sum of partions. check for negative variances")}
```

```
  return(r2.res)}
```
